# Genomic and Genetic Insights into Mendel’s Pea Genes

**DOI:** 10.1101/2024.05.31.596837

**Authors:** Cong Feng, Baizhi Chen, Julie Hofer, Yan Shi, Mei Jiang, Bo Song, Hong Cheng, Lu Lu, Luyao Wang, Alex Howard, Abdel Bendahmane, Anissa Fouchal, Carol Moreau, Chie Sawada, Christine LeSignor, Eleni Vikeli, Georgios Tsanakas, Hang Zhao, Jitender Cheema, J. Elaine Barclay, Liz Sayers, Luzie Wingen, Marielle Vigouroux, Martin Vickers, Mike Ambrose, Marion Dalmais, Paola Higuera-Poveda, Rebecca Spanner, Richard Horler, Roland Wouters, Smitha Chundakkad, Xiaoxiao Zhao, Xiuli Li, Yuchen Sun, Zejian Huang, Xing Wang Deng, Burkhard Steuernagel, Claire Domoney, Noel Ellis, Noam Chayut, Shifeng Cheng

**Author notes:** These authors contributed equally to this article (Cong Feng, Baizhi Chen, Julie Hofer, Yan Shi). These authors contributed equally to this article (Mei Jiang, Bo Song, Hong Cheng).

## Abstract

Pea, *Pisum sativum*, is an excellent model system through which Gregor Mendel established the foundational principles of inheritance. Surprisingly, till today, the molecular nature of the genetic differences underlying the seven pairs of contrasting traits that Mendel studied in detail remains partially understood. Here, we present a genomic and phenotypic variation map, coupled with haplotype-phenotype association analyses across a wide range of traits in a global *Pisum* diversity panel. We focus on a genomics-enabled genetic dissection of each of the seven traits Mendel studied, revealing many previously undescribed alleles for the four characterized genes, *R*, *Le*, *I* and *A*, and elucidating the gene identities and mutations for the remaining three uncharacterized traits. Notably, we identify: (1) a ca. 100kb deletion upstream of the *Chlorophyll synthase* (*ChlG*) gene, which generates aberrant transcripts and confers the yellow pod phenotype of *gp* mutants; (2) an in-frame premature stop codon mutation in a Dodeca-CLE41/44 signalling peptide which explains the parchmentless mutant phenotype corresponding to *p*; and (3) a 5bp in-frame deletion in a *CIK-like* receptor kinase gene corresponding to the fasciated stem phenotype *fa*, which Mendel described in terms of flower position, and we postulate the existence of a *Modifier of fa* (*Mfa*) locus that masks this meristem defect. Mendel noted the pleiotropy of the *a* mutation, including inhibition of axil ring anthocyanin pigmentation, a trait we found to be controlled by allelic variants of the gene *D* within an *R2R3-MYB* gene cluster. Furthermore, we characterize and validate natural variation of a quantitative genetic locus governing both pod width and seed weight, characters that Mendel deemed were not sufficiently demarcated for his analyses. This study establishes a cornerstone for fundamental research, education in biology and genetics, and pea breeding practices.

## MAIN TEXT

Pea is an Old World crop first brought into cultivation about 10,000 years ago in the Fertile Crescent^1^. Pea is mainly grown as a field crop, with about ¾ of the area for dry seed and ¼ for use as a vegetable, totalling about three billion USD in export value in 2022 (https://www.fao.org/faostat/en/#data/). Pea also has a minor use as a fodder crop and is often grown in home gardens. The nutritional and environmental benefits of this pulse crop have been discussed elsewhere^2,3^.

Pea has considerable diversity, both genetically and phenotypically. The nucleotide diversity in *Pisum* (from π = 8.2 × 10^−4^ among wild *Pisum* to π = 2.4 × 10^−4^ in cultivars)^4^, is about tenfold greater than that in the human population^5^, reflecting bidirectional introgression between the cultigen and wild genotypes. *Pisum sativum* (meaning cultivated pea) is a subset of *Pisum* as a whole; wild peas designated *P. fulvum* are noticeably distinct^6,7^ and carry a translocation with respect to the rest of *Pisum*^4^, which creates a fertility barrier. Similarly, the independently domesticated *P. abyssinicum*^8,9^ differs in karyotype with respect to the rest of *Pisum*^4^, again presenting a fertility barrier. Thus, *Pisum* comprises four major divisions; the cultivated forms *P. sativum* and *P. abyssinicum* and the wild forms *P. fulvum* and *P. elatius*.

The morphological diversity within *Pisum* has been documented since at least the 16^th^ century, with Gerard^10^ (p1045) illustrating four forms: *P. majus, P. minus, P. umbellatum* (fasciated) and *P.excorticatum* (parchmentless) and discussing several others, such as those with seeds “which being drie are cornered”. If by this description, Gerard was referring to wrinkled peas, then three of the variant forms that Mendel studied^11^, viz. peas with stem fasciation, parchmentless pods, and wrinkled seeds, had been recorded nearly 300 years earlier, while the white flowered forms, as previously noted, were described about 1300^12^. Pea is predominantly inbreeding, with large flowers; these two features, and the many easily distinguishable characteristics of pea, made this species ideal for Gregor Mendel’s studies of inheritance using hybridization^11,13^. For example, the seven variants that Mendel studied in detail were clearly distinguished in the seed catalogues of the time^14^, representing different agronomic forms, end uses, or market types, as they still do today.

Mendel’s work on peas was described by Allan Franklin as “*The best experiments ever done*”^15^. Pea serves as an excellent plant model system; in addition to its significant historical contribution to the development of genetics, approximately 60 pea genes have been characterized at the molecular level^16^. However, much remains unknown about the molecular nature of the contrasting traits that Mendel studied, even though the genetic loci were named over a century ago^17^. The four cloned genes *R*, *Le*, *I* and *A* have been characterized for some time^12,18–23^, but the extent of their natural allelic variation, its distribution and genomic context is still largely unknown^4,16,24^. The gene identities of the remaining three Mendel traits, *P* (or *V*, pod form), *Gp* (pod colour) and *Fa* (or *Fas*, fasciation), remain uncharacterized. Candidates for *Gp* and *P* have been tentatively proposed, based on specific GWAS analyses and bi-parental mapping studies^25,26^; however, further work is needed to confirm or reject these proposals.

In this study, we couple sequence-based genomic diversity analysis with phenotypic variation to elucidate gene identity underlying traits of interest in one of the world’s major *Pisum* germplasm collections^16^. We illustrate this by describing the genomic context of the seven well-known traits that Mendel studied in detail. We further demonstrate how this can be expanded to elucidate the molecular basis of other characters, including several quantitative traits that Mendel discussed but considered too variable for simple analysis.

## RESULTS

### Genomic Variation Map of a *Pisum* Core Collection

To build a pea genomic variation map, and particularly to characterize each of the genetic loci underpinning the traits that Mendel studied, we selected a core diversity panel from the JI *Pisum* Germplasm Collection, a widely-used collection, historically and globally^16^. The panel included 500 representative *Pisum* accessions, selected using Corehunter 3 and based on prior genotyping data^27,28^. This set was augmented by the inclusion of an additional 130 lines previously chosen for other diversity studies (www.pcgin.org) and included parents of mutant and mapping populations together with 67 lines comprising all accessions designated *P. abyssinicum, P. humile* or *P. fulvum* (Fig. 1a, Supplementary Table 1). We performed next-generation short-read whole-genome resequencing for these 697 *Pisum* accessions, resulting in approximately 80 Gb of clean reads, with a coverage of about 20X for each accession (Supplementary Table 2). We built a genomic variation map encompassing 154.8 million high-quality single nucleotide polymorphisms (SNPs) with respect to the ZW6 assembly^24^, as well to Caméor v1a^4^ (Supplementary Table 3-4). This revealed the pattern of accession relationships and defined population structure at a high resolution within *Pisum* (Supplementary Table 5) which is broadly consistent with previous results^28^, and we proposed eight major *Pisum* groups (G1-G8) (Fig. 1b, 1d-e). These accessions do not have a tree-like relationship but have a reticulated network structure (Fig. 1e). Within the diversity panel, we particularly recorded the phenotypic variation for each of the seven pairs of contrasting traits that Mendel studied (Fig. 1c) and associated this with genomic diversity (Fig. 1f and Supplementary Table 6-11).

**Fig. 1.**
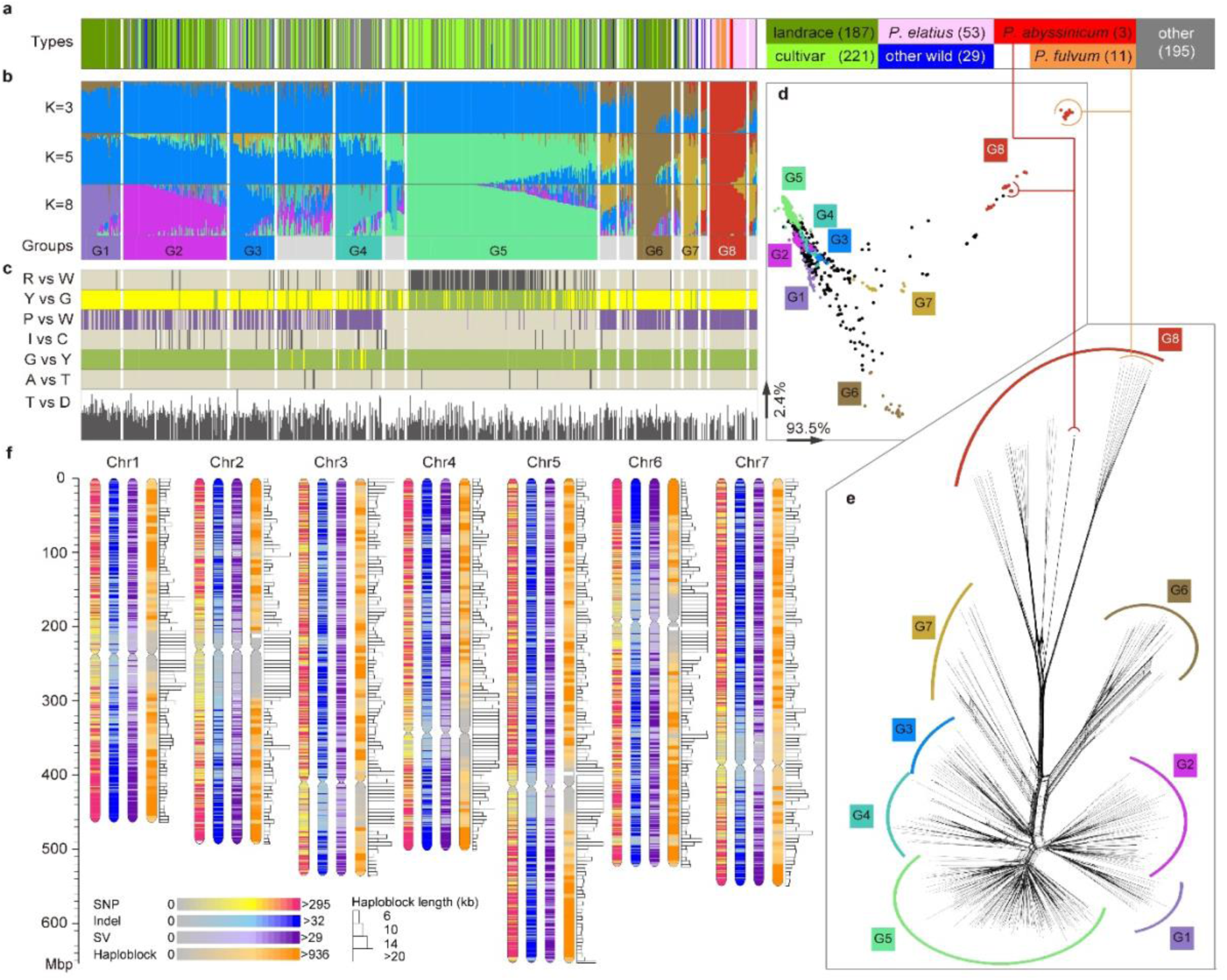
Genotypic and phenotypic variation with respect to population and genome structure within *Pisum*. **a,** Taxa types and other classifications as indicated by colour on the right, including the wild taxa: *P. fulvum*, *P. elatius*, and ‘other wild’ (various named taxa Supplementary Tables 1, 5) and domesticated taxa: *P. abyssinicum* and *P. sativum* classified into ‘cultivars’, ‘landraces’, and ‘other’ that mostly comprises genetic stocks. The number in the brackets indicates the number of accessions for each classification. **b,** Admixture K = 3 (average of 5 runs), Admixture K = 5 (average of 3 runs), Admixture K = 8 (one run that splits K = 5 groups) and accessions strongly assigned to Admixture groups (by colour, grey = admixture) (corresponding to Supplementary Table 5); **c,** Distribution of phenotypes in Mendel’s seven pea traits, with initials labelled: R (Round, pale) vs W (Wrinkled, black), seed shape; Y (Yellow) vs G (Green), cotyledon colour; P (Pigmented, purple) vs W (White, pale), flower colour; I (Inflated, pale) vs C (Constricted, black), pod shape; G (Green) vs Y (Yellow), pod colour; A (Axial, pale) vs T (Terminal, black), flower position; and T (Tall) vs D (Dwarf), internode length. The bar is proportional to internode length. **d,** PCA of PLINK distance matrix for all accessions, those with Q-value >0.75 indicated by colour. **e,** Splits Tree^81^ analysis of accessions with Q-value >0.75 indicated by colour. **f**, *Pisum* genomic variation map distributed along all seven chromosomes, including SNPs, insertions and deletions (<50bp), large-scale structural variations (SV), and the linkage disequilibrium (LD) based haplotype map.

### Novel Alleles for Mendel’s Four Characterized Genes

Haplotype-phenotype association coupled with linkage analysis of bi-parental mapping populations elucidated the genetic basis of Mendel’s pea traits and revealed their genetic structure (Fig. 2). From the significance of the association between SNP variants and the phenotypic differences which Mendel described^11,13^, we can see that for each trait, a small number of specific genetic loci contribute to the trait variation. Our novel discoveries are summarized (Extended Data Fig. 1) and explained below for each trait.

**Fig. 2.**
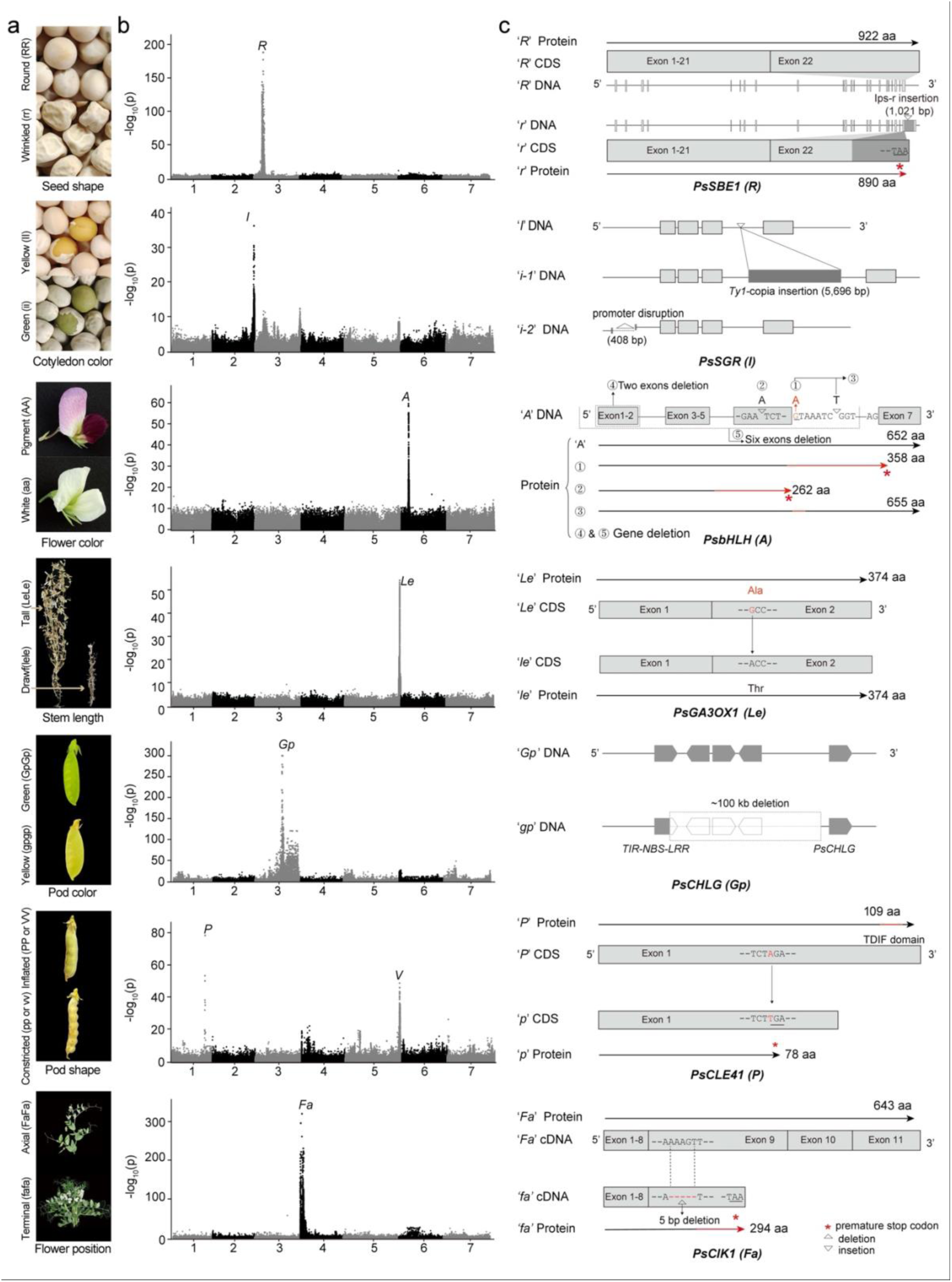
Genetic architecture and genomic diversity of the genes underlying the seven traits that Mendel studied. **a**, Pictures of the contrasting phenotypes of the seven traits. **b**, Manhattan plots from the whole genome-wide association study (GWAS) for phenotypic differences of each trait as scored in this study and plotted against the ZW6 assembly. **c**, Gene models for wild type and natural mutant alleles for each of the seven traits (more details are described in the Main Text, Extended Data Figures and Supplementary notes).

#### Round *vs* wrinkled seeds

Our association genomics analysis of round and wrinkled seeds (Supplementary Table 12) identified a single strong but broad signal, at the expected genomic position of *R*, encoding Starch Branching Enzyme I^23^(*PsSBE1*). The insertion of *Ips-r*, a 1021bp non-autonomous *Ac-* like transposable element, within exon 22 of the *PsSBE1* coding sequence, predicts a truncated protein (from 922 aa to 890 aa) due to a premature stop codon^23,29^ (Fig. 2c, Extended Data Fig. 2), although multiple *r* transcripts have also been detected^30^. Genetic differentiation between round and wrinkled types in breeding programmes could be the underlying reason for the broad GWAS peak: round types are field peas, grown for their dry seed, while wrinkled types are a class of peas grown for harvesting before maturity as fresh peas or for the freezing market^31^. The single GWAS peak also indicates that there is no genetic heterogeneity associated with this phenotype within the set of lines we have examined.

#### Green *vs* yellow cotyledons

We identified a strong signal at the expected position of *I*, the gene encoding Mg-dechelatase^20–22^ which catalyses the first step in chlorophyll degradation and underlies the genetic difference between green vs yellow cotyledons (Supplementary Fig. 1, Supplementary Table 13). We found two classes of *i* alleles (Fig. 2c and Extended Data Fig. 3), which explain most of the green cotyledon mutants in this diversity panel. The more common mutant allele (designated as *i-1*) is the insertion of a 5,696 nt TAR element (a *Ty1*-*Copia* LTR retrotransposon) which probably corresponds to group 3 alleles as previously suggested^21^ but not identified, nor was its frequency characterized at the population level. The second allele we discovered is a novel 408 bp deletion in the promoter of the Mg-dechelatase gene (Supplementary Fig. 2), which we designate as the ‘*i-2*’ allele, explaining 15 accessions with green cotyledons. Neither the 6bp insertion event corresponding to the *i^JI^*^2775^ allele^21^ nor the *i^PI^* allele^21^ described as group 4 was found in any accession of our diversity panel (JI2775 was not included in this study), so it is presumed that these alleles are rare. Several minor peaks with a -log_10_(p) value ∼10, distinct from the *I* locus, can be seen in the Manhattan plot (Fig. 2b). Modifiers of cotyledon colour are well known and ten genetic loci that contribute to this effect have been identified^32^, but their location with respect to these additional GWAS peaks could not been determined.

#### Presence or absence of anthocyanin pigment

Our haplotype-phenotype association study of pigmented vs white flowers (Supplementary Table 14) revealed a single strong signal spanning a genomic region consistent with the location of *A*, which encodes a *bHLH* transcription factor^12^ (Fig. 2) required for the expression of chalcone synthase in epidermal tissues^33^, thereby enabling anthocyanin pigmentation. We discovered several novel haplotypes within the structural gene (Extended Data Fig. 4). The wild types (*A*) with pigmented flowers were assigned to haplotype (Hap) 1 based on the distribution of functional variants, but Hap1 is remarkably diverse at other positions. The two most common *a* alleles correspond to Hap5, carrying the splice donor site variant (G to A), originally identified in Caméor, and Hap2, with an additional ‘A’ nucleotide in exon 6 creating a premature stop codon, as originally identified in JI1987^12^. Two new variants (Hap3, with four accessions and Hap4, with one accession) are deletions of part (the first two exons), or almost the entirety (the first six exons) of the gene.

Remarkably, we found one accession with coloured flowers that carried the splice donor site mutation which should render the gene dysfunctional. This allele (in JI0233, Hap5) has an additional ‘T’ nucleotide in what would be the sixth intron of the wild type allele, but it lies between the wild-type splice donor site and the splice site used in the Caméor allele, adding nine nucleotides to the transcript, one more than in the *a^Caméor^* allele (Extended Data Fig. 4). Thus this ‘T’ insertion is an intragenic suppressor mutation, which restores wild type gene function by restoring the reading frame in the JI0233 transcript, resulting in the predicted addition of three amino acids to the A protein (Supplementary Fig. 3).

#### Internode length

Variation in internode length in our analysis corresponds to Mendel’s plant height character (Supplementary Table 15). We identified a significant peak (chr5: 620824850-652929960) at the end of chromosome 5, which spans the location of *Le* encoding GA 3-oxidase1 (Psat05G0825300, also called GA 3β-hydroxylase) (Fig. 2), but does not extend to *Lh*^34^ (Psat05G0840800, chr5:650785676-650788204), another gene conditioning plant height, closely linked to *Le*. That the GWAS approach finds this single peak suggests that variation at other known loci involved in regulation of the type and abundance of plant hormones affecting plant height, or internode length, does not contribute significantly to natural phenotypic variation in this trait. We observed five haplotypes associated with *Le* (Extended Data Fig. 5), but the reduced height *le* variants were exclusively found in haplotype 1, which carries the known G-A substitution at chr5:639901919^18,19^.

### GENE IDENTITY AND VARIATION OF THE UNCHARACTERIZED TRAITS

Three of Mendel’s seven traits have remained poorly characterised^35^: ‘the difference in the colour of the unripe pod’ (*Gp*), ‘the difference in the shape of the ripe pod’ (conditioned by either of two loci, *P* or *V*), and ‘the difference in the position of the flowers’ (thought to be conditioned by either of two loci, *Fa* or *Fas*). Gene identities and allelic variation underlying these traits were investigated in this study.

#### Pod colour

Although *Gp* is usually discussed in relation to pod colour, Mendel noted that yellow pods are just one feature of the *gp* mutant. In mature flowering and fruiting plants, yellow tissues are seen in the petiole, rachis, tendrils and leaflet midribs of young leaves, and also in the pedicel, peduncle and sepals (Fig. 3a and Supplementary Fig. 4). There are also significant differences in the physiological and biochemical properties of pod and leaf tissue, and differences in chloroplast development^36^, between the green (*GpGp*) and the yellow podded (*gpgp*) varieties (Fig. 3b). Here we found that even the green leaves of *gp* lines have disturbed development of thylakoid membranes (Fig. 3c) and this was reflected in a difference in productivity between *Gp* and *gp* isolines (Supplementary Fig. 5).

**Fig. 3.**
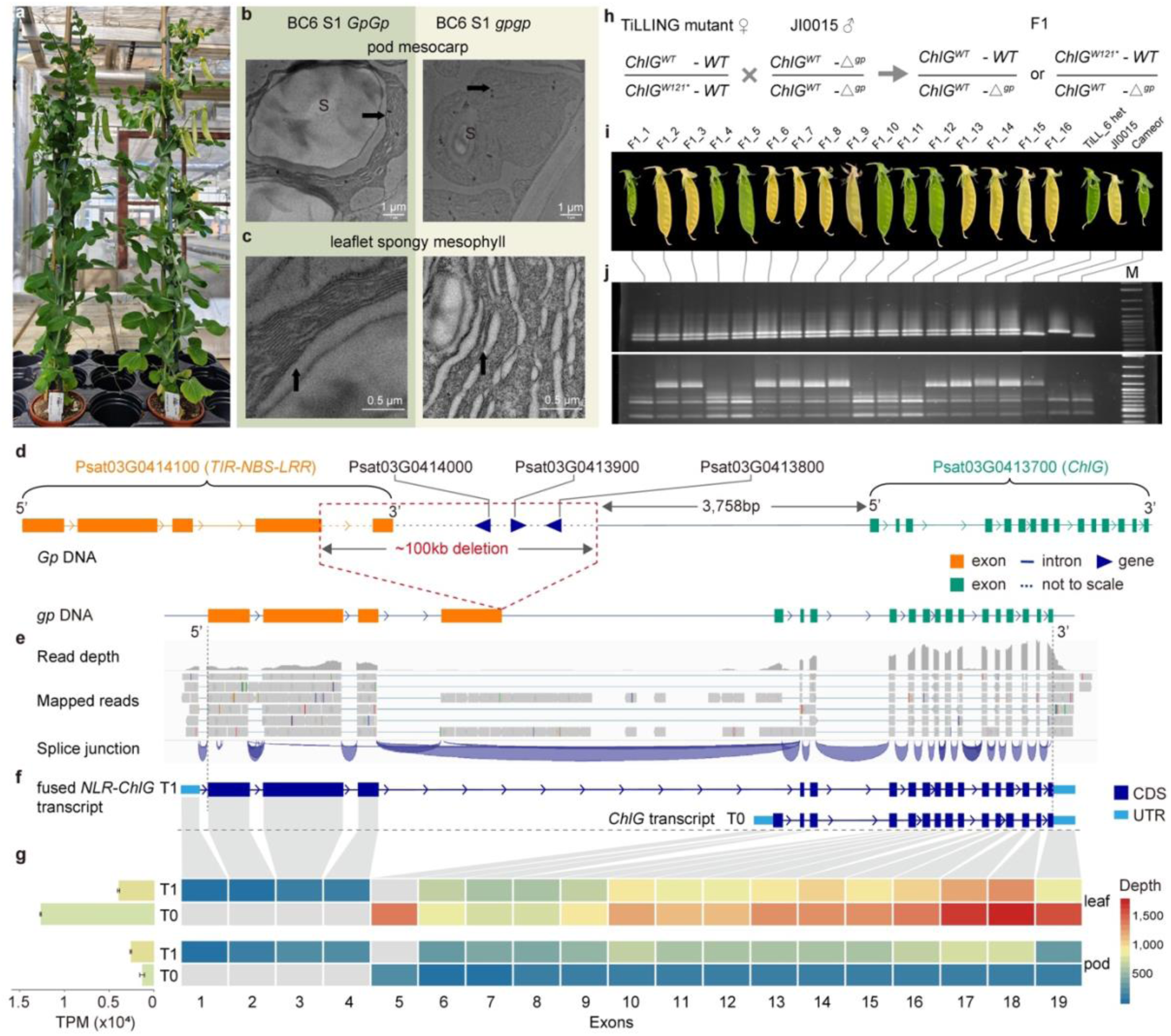
The *gp* mutant. **a**, General view of near-isogenic plants (BC6 S1of the cross JI0015 *gpgp* x Caméor *GpGp*) developed in this study. Pot size is 9 cm diameter. Note the yellow peduncle and pale sepals as well as the yellow vs green pods on *gp* compared to *Gp*. **b**, TEM sections of pod mesocarp cells. Note the large starch grain (S) in *Gp* compared to *gp* and the poorly developed thylakoid membranes (arrows) in *gp* compared to *Gp*. **c**, TEM sections of leaflet spongy mesophyll cells. Note the poorly developed thylakoid membranes (arrows) in *gp* compared to *Gp*. **d**, A ca. 100 kb deletion adjacent to *ChlG* is illustrated for *gp* compared to ZW6 (*Gp*), the deletion event in *gp* lines is illustrated on a *Gp* reference genome by dashed box, affecting five genes. The ca. 100 kb deletion event was called using ZW6 as the reference, the gene content and orientation is based on ZW6 genome annotation. **e**, RNA-seq data shows the pattern of read-through transcription in *gp* lines (JI0015 and JI2366) across the ca. 100kb deletion and internally within *ChlG*. **f**, Predicted structure of two transcripts in *gp* mutant pods: a fused aberrant transcript (T1) and the *ChlG* transcript (T0). **g**, RPKM counts for exons adjacent to the ca. 100 kb deletion compared between leaves and pods of JI2366; the horizontal bars in the left indicate the average transcript abundance (measured by TPM) of T1 and T0 in both pods and leaves. **h**, Crossing scheme for a complementation test between Caméor M4 TILLING line 411.1 carrying one lethal allele of *ChlG* and *gp* (JI0015), with the two types of expected F1 genotype. *ChlG^WT^* and *ChlG^W^*^121^*\** represent the wild type and TILLING mutant of *ChlG. WT* represents the presence of the wild type (Caméor) sequence between *ChlG* and the *TIR-NBS-LRR* gene, while *Δ^gp^* represents the ca. 100 kb deletion which co-segregates with *gp*. The question being addressed is whether *ChlG^W121*^ - WT* does or does not complement *gp* (*ChlG^WT^ – Δ^gp^*). **i**, F1 pods segregating for green vs yellow, F1_x – indicates plant number; the parental lines (TILL_6 het and JI0015) and wild type Caméor are also indicated. **j**, Codominant PCR marker test confirming all plants presumed to be F1s are *Gpgp* heterozygotes (upper panel) and a dCAPS marker PCR test confirming that only the yellow podded F1 plants inherited the *ChlG^W121*^* TILLING allele. M, DNA size marker.

All yellow podded lines in the JI *Pisum* germplasm collection were shown to be allelic to *gp*. Thus, there is only one known yellow pod locus and we show below that there is only one yellow podded *gp* allele. Genetic mapping and association genomics analysis found that all these yellow podded lines carried a ca. 100 kb deletion within the GWAS interval, which co-segregated with *gp* (Supplementary Figs. 6-8 and Supplementary Table 16-21). With respect to the ZW6 assembly, this deletion removes three entire genes, as well as part of exon5 and the whole of exon 6 from a gene encoding a *TIR-NBS-LRR* (*NLR*, *Past03G0414100*) protein.

Interestingly, this deletion is adjacent to the gene encoding chlorophyll synthase (*ChlG, Psat03G0413700*), but the structure of the *ChlG* gene is intact in all the *gp* lines and the encoded amino acid sequence is identical to the wild type (Fig. 3d). Mapping RNA-seq reads to their matched *gp* genome assemblies of JI2366 and JI0015 predicted novel transcripts from *ChlG*, including intron read-through and a fusion of the truncated *TIR-NBS-LRR* and *ChlG* transcripts, confirmed by transcriptome sequencing (Fig. 3e-f, Supplementary Figs. 9-10). Furthermore, RNA-seq and qPCR data showed that *ChlG* transcript abundance was reduced in *gp* pods with respect to *gp* leaves, whereas the abundance of the fused *NLR-ChlG* transcript in *gp* lines is similar in pods and leaves (Fig. 3g). We propose that disruption of chlorophyll synthesis by transcriptional interference from the expression of aberrant transcripts is the reason for the yellowing of otherwise green tissues in the *gp* mutant (Supplementary Note).

To test the hypothesis that *Gp* corresponds to *ChlG*, we obtained a TILLING mutant^37^ with a premature stop codon (W121*) in *ChlG* (Fig. 3h). This mutant could not be recovered as a homozygote, although the mutant allele could be transmitted through both pollen and egg cells, so we conclude that the homozygous mutation is embryo lethal, but it is not lethal in either gametophyte. We reasoned that the phenotype of a *Gpgp*, *ChlG^wt^ChlG^W^*^121^*\** double heterozygote would be informative; if *Gp* did not correspond to *ChlG* then it should be viable and green-podded and our hypothesis would be refuted. Conversely, if *Gp* did correspond to a functional *ChlG* then it should be yellow podded. Of the sixteen F1 double heterozygotes we derived from the cross between *gpgp* and the TILLING mutant heterozygotes, half had yellow pods, and all of these yellow podded F1s carried the *ChlG^W1^*^21^*\** null allele (Fig. 3i-j). This result upheld our hypothesis and showed that the *gp* mutant does not provide a fully functional *ChlG*.

The evidence presented above demonstrates that a *ChlG* deficiency mediates the mutant phenotype, and establishes that *ChlG* is allelic to *Gp*. The large genomic deletion upstream of *ChlG* in the *gp* lines generates fused aberrant transcripts spanning *ChlG* and an upstream *TIR-NBS-LRR* gene. The detailed molecular mechanism of this defect in chlorophyll synthesis and the possible role of the other genes affected by the deletion event remain to be established; however, our current understanding predicts that ablation of the *NLR* gene in a *gp* mutant, thereby removing the fused *NLR-ChlG* transcripts, would restore the wild-type green pod colour.

#### Pod shape

The difference in the shape of the ripe pod was described by Ruel in 1537 as ‘Valvulae etia recetes eorum quae nullo pedameto fulciuntur, ante que durescat, edendo sunt’^38^ which roughly translated means ‘Those where the valves provide little support are to be eaten before they harden’, indicating that, as today, these are a vegetable form. The lack of a sclerenchyma layer in pea pods (pod parchment) is conditioned by the recessive allele at either (or both) of the genes *P* and *V*. It is uncertain which of these genes Mendel was discussing; he could have worked with either, or perhaps both (Supplementary Note). Mendel used this parchmentless variant in several crosses, including the four factor cross described in his second letter to Nägeli^39^. Our GWAS analysis identified several regions that are statistically correlated with this phenotype (Supplementary Table 22) and of these, two correspond to the expected positions of *P* and *V* (Fig. 2b), suggesting that both *p* and *v* alleles are relatively common. The additional signals may correspond to genes affecting pod wall thickness (*N*) or structure (*Sin*)^40^ (Extended Data Fig. 6)

Notably, within our 8.3Mb GWAS peak at the end of Chr1 (Fig. 2b), the gene *Psat01G0420500* had the greatest significance, which is consistent with a 0.92Mb interval defined in the JI0816 x JI2822 F2 mapping population (Supplementary Table 17-19, Extended Data Fig. 6a-d). *Psat01G0420500* is annotated as encoding a Dodeca-CLE peptide and includes the tracheary element differentiation inhibitory factor (TDIF) of CLE41/44^41^. One allele of this gene, carrying an in-frame premature stop codon (R79*) upstream of the TDIF motif (Fig. 2c, Extended Data Fig. 6e-g), fully explains the *p* phenotype. CLE41 peptides repress the formation of xylem^42^ and specify positional information that determines the rate and orientation of cell divisions in vascular tissue in conjunction with the receptor kinase PXY39^43^. TDIF is proposed to be a non-cell autonomous signalling peptide controlling cell fate^44^ and lignification^45^. This suggests a model for *P* whereby this TDIF peptide interacts with a PXY-like protein to specify pea pod sclerenchyma development.

The genomic interval corresponding to *V*, as identified by GWAS, spans a broad region (Chr6 610-650Mb). A 3Mb interval (Chr6 628-631Mb) in the middle of the GWAS peak, was the most significant location for the identification of candidate genes to *V* (Extended Data Fig. 7, Supplementary Table 23). Within this interval, we found that accessions with parchmentless pods, including those which lack the R79* mutation in CLE41/44 (*PPvv*) and those with the double mutantion (*ppvv*), are clustered into haplotype 2 of *Psat05G0805200,* a cell wall invertase. While this gene is a plausible candidate for *V* further work is needed to fully explain the *v* alleles (Supplementary Note and Extended Data Fig. 7 and Supplementary Fig. 12).

#### Fasciation

Mendel discussed “the position of the flowers” on the stem of pea and used the name *Pisum umbellatum*, a term previously used by Gerard^10^ to describe the fasciated form (Supplementary Table 24) with an umbellate inflorescence. In pea, fasciation can vary in its severity, from stem bifurcation to an extreme clustering of flowers at the apex. We conducted a comparative analysis of field phenotypes and microscopic observations in the apical meristem of fasciated vs wild type plants (Supplementary Fig. 13). The bunched apical flowers of the mutant are borne on a wider stem with additional vascular strands derived from a broadened apical meristem. There are several pea genes, which when mutant, have a fasciated phenotype; of these, *Fa* vs *fa* (chromosome 4 linkage group IV) is considered to be the gene Mendel studied^46,47^.

Our GWAS analysis identified a broad signal (Chr4 0-40Mb) (Fig. 2b), which underwent further refinement through investigation of F2 populations using bulked segregant analysis (BSA), narrowing the interval down to a 15Mb region (Supplementary Fig. 14). Subsequent fine-mapping led to the delineation of a 1.33Mb interval (Chr4 18.18-19.51Mb, ZW6) (Extended Data Fig. 8a-e, and Supplementary Table 17-19, 25-26). We found that all the accessions with fasciated phenotypes were clustered together within haplotype 5 of this 1.33Mb interval (Extended Data Fig. 8f); however, JI1713 and JI0815 in haplotype 5 are not fasciated (see explanation below and in the supplementary notes). A similar analysis was performed with each gene within this interval, revealing a significant finding: only one gene, *Psat04G0031700*, co-segregated with fasciation. All accessions with the recessive phenotype (fasciation) are clustered into haplotype (Hap) 3 of this gene, which is characterized by a 5bp deletion in exon 2, creating a frameshift and premature stop codon which would render the protein non-functional, thereby explaining fasciation in *fa* lines (Extended Data Fig. 8g, h). This gene encodes a cell membrane-localized Senescence-Associated Receptor-Like Kinase, a class of *CLAVATA3 INSENSITIVE RECEPTOR KINASES* (*CIK*) signalling receptor kinases known for their role in maintaining the structure of the shoot apical meristem^48^. Our hypothesis that a module involving *PsCIK*, identified here, and *PsWUS* and *PsCLV3* (Supplementary Fig. 15), key genes expressed in the shoot apex and known to be involved in meristem maintenance in other contexts^49^, can now be tested using biochemical genetics.

There is a second unexpected minor signal on chromosome 6 linkage group II in our GWAS analysis, which is consistent with the BSA analysis showing a small signal at chr6LGII (Supplementary Notes). In the JI0816 (*fa*) x JI2822 (*Fa*) F2 population (Supplementary Table 17-19), we noticed that out of 395 scored individuals, 32 had a wild-type phenotype but carried the recessive allele at *fa* (Extended Data Fig. 9), as was also the case in the GWAS and BSA studies. This suggests a model whereby the recessive allele of a gene in this region at chr6LGII masks the fasciated phenotype. Accordingly, we designated this second locus as “*modifier of fa”* (*mfa*) (Supplementary Notes). In this model, individuals that are recessive for both loci, the *fafa mfamfa* genotype (double recessive), have a wild-type appearance. This proposal would explain why some accessions, like JI1713 and JI0815, carry the 5bp deletion in *Psat04G0031700* (*PsCIK1*) but are not fasciated. Previous studies have highlighted complexity in the segregation of fasciation, with reports of both reversals of dominance and two-factor segregation ratios (15:1) in F2 populations for some crosses^50^, rather than the expected one-factor segregation ratio (3:1). These unusual features may, in part, be explained by the previously unrecognised gene *Mfa* (Extended Data Fig. 9). The nature of *Mfa* remains to be determined, but it resides within the interval ZW6 Chr6: 244,689,457-253,701,016 identified in this study (Supplementary Fig. 14).

### From Mendel’s Genetic Loci to Quantitative Traits

It has been argued that Mendel’s motivation in studying inheritance was related to an applied plant breeding program^51^. In this work, we measured 74 additional agriculturally relevant characters within our *Pisum* diversity panel, including seed, pod, flower, leaf, and plant architecture traits (Supplementary Table 27, Extended Data Fig. 10a,b). A comprehensive genome-wide association study established hundreds of significant marker–trait associations (Supplementary Table 28), including several previously cloned loci such as *Er1*^52^*, Pl*^53^, *Af*^54^*, Tl*^55^*, Rms1*^56^*, Hr*^57^*, St*^58^*, Rms3*^59^*, K*^60^*, Rms4*^59^ and *Sn*^61^ (Fig. 4a, Supplementary Table 29). In addition, our analyses clearly determined the physical locations of 20 historically defined genetic loci (Fig. 4a), to within an average genomic interval 12 Mb (ca. 150 protein-coding genes). Examples include: the *Aero* locus (at the end of Chr2), associated with silver flecking on pea stipules^62^; the *Bt* locus (at the beginning of Chr3), influencing the pointed tip of the pod^63^; and the *N* locus (at the beginning of Chr4), enhancing pod thickness for snap peas^40^. Furthermore, in addition to the three newly characterized of Mendel’s pea traits, our study uncovered several potentially important new loci: the *LC* (Leaf Colour) locus on Chr1, impacting leaf colour intensity, and Organ size locus (*Os1*), controlling pod width and grain weight which was validated below. These results demonstrate the high-quality of our dataset and the reliability of the association genomics analyses, laying a solid foundation for future functional elucidation in peas, both for fundamental research and pea breeding.

**Fig. 4.**
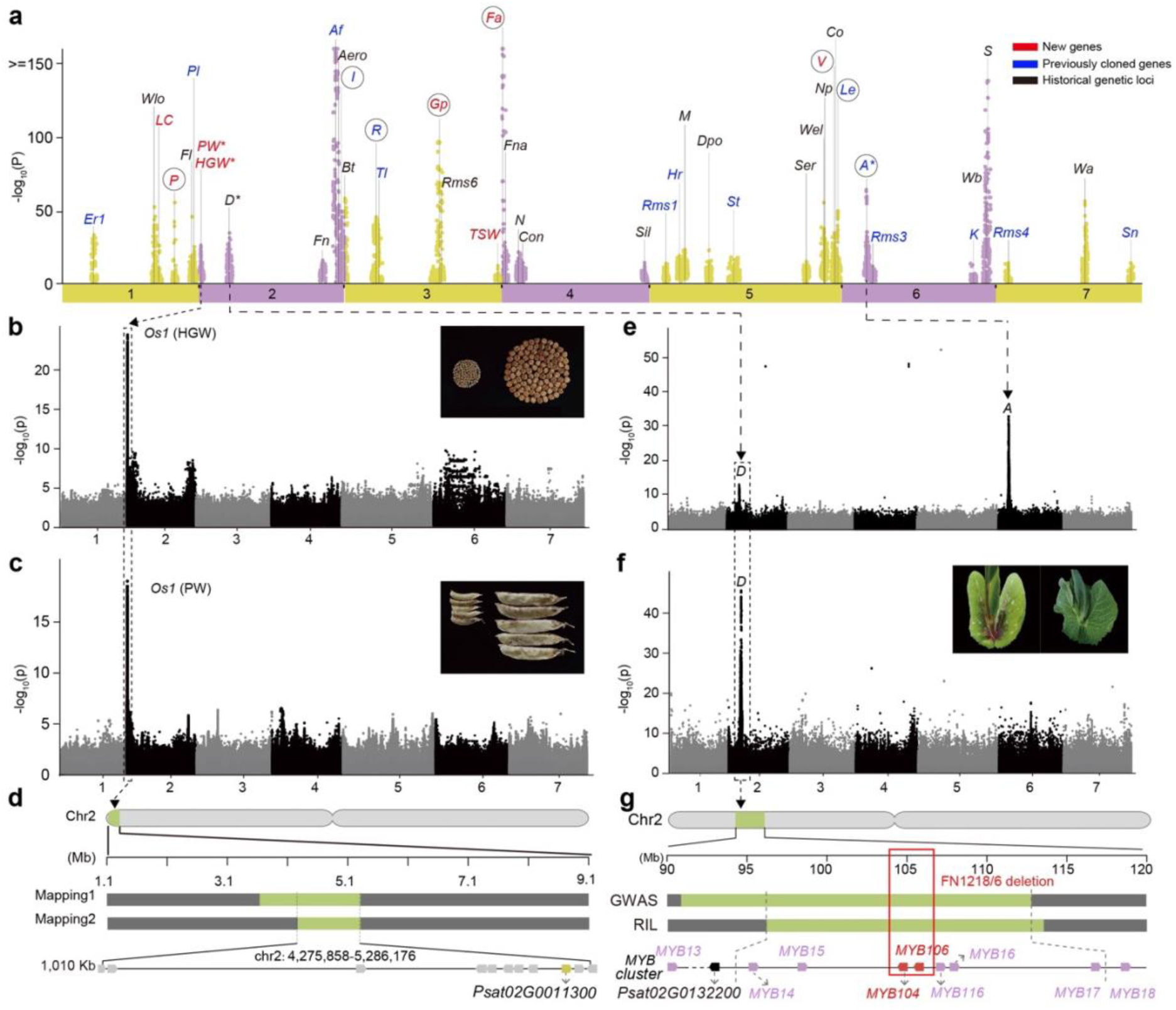
A Genome-Phenome association map for identification of genetic loci that confer agronomic traits. **a,** Summary of the most significant trait-marker associations underlying a variety of agronomic traits presented as a combined Manhattan plot. The detailed information for each of these genetic loci is in Supplementary Table 28. Gene symbols marked in a circle correspond to Mendel’s loci; symbols in red indicate novel genetic loci or genes discovered in this study; symbols in blue are previously characterised and cloned genes positioned with respect to the ZW6 assembly; symbols in black are suggestions from known genetic map locations, but without specific gene candidate and position. **b**, Manhattan plot of GWAS data relating to hundred grain weight (HGW). **c**, Manhattan plot of GWAS data relating to pod width (PW). The HGW and PW genomic intervals span the same 8Mb genomic region, named *Organ Size 1* (*PsOs1*). **d**, Narrowed genomic interval of *PsOs1* on Chr2 defined by two F2 mapping populations and BSA analysis (Supplementary Fig. 23-24, Online Method) as a 1.01Mb region encompassing 11 protein-coding genes, in which *Psat02G0011300* (marked in yellow) is the most highly expressed gene. *Psat02G0011300* encodes a SIAMESE-related protein (SIM/SMR), a cyclin-dependent protein kinase inhibitor, the gene functional validation was presented in supplementary note and supplementary Figs. 25-30. **e**, Manhattan plot of GWAS data on the presence or absence of axial ring pigmentation across our diversity panel, using phenotypic data collected at Shenzhen (2021); **f**, Manhattan plot of GWAS data on the presence or absence of axial ring pigmentation, on a subset of phenotypic data excluding accessions carrying the white flowers (*aa*). These data were collected at Harbin (northern China, 2022). A peak at the expected genomic position of *D* is significantly associated with the accumulation of axillary anthocyanin, and the peak at Chr6 is the location of *A*. **g**, Genomic interval of *D* on Chr2 defined by RIL mapping, GWAS analyses, further defined by bioinformatic analysis of Fast Neutron mutants as a *MYB* gene cluster^12,64,65^, with the genes *PsMYB104* and *PsMYB106* both deleted in the *d* mutant line, FN1218/6. The box outlined with a dashed line indicates the approximate position of the deletion detected in FN1218/6 from mapping of sequence reads. Inset photographs show the contrasting phenotypes in every case.

#### Genetic complexity that Mendel discussed: pleiotropy and epistasis

In his 1866 paper, Mendel noted the pleiotropic effects of the seed coat/flower colour trait (*A vs a*) and specifically referred to the presence or absence of axil ring pigmentation as one of these effects. *A* regulates the presence or absence of anthocyanin pigmentation throughout the plant and *a* is epistatic to *d*, which regulates the pattern of axil ring pigmentation^50^. The range of axil pigmentation patterns in pea (Supplementary Fig. 16) is reminiscent of leaf marking in *Trifolium*^64^ and *Medicago*^65^ both of which are controlled by similar *MYB* transcription factors.

Our GWAS analysis revealed two strong signals associated with axil ring pigmentation (in coloured flower lines) (Fig. 4e-f, Supplementary Table 30). One of these corresponds to *A* (Chr6), while the other is at the expected position of *D* (Chr2) where there is a cluster of *MYB* genes^66^ (Supplementary Fig. 17). The potential role of one of these *MYB* genes was investigated further by Virus-Induced Gene Silencing (VIGS), which showed that the *MYB*-encoding gene *Psat02G0138300* (*PsMYB16*) affect the accumulation of the axil ring anthocyanin pigmentation (Supplementary Figs. 18-19). Furthermore, deletion of another two *MYB* genes at the same locus, *PsMYB104* and *PsMYB106*^67^, in the Fast Neutron (FN) induced mutant line FN1218/6, resulted in the complete absence of axil ring pigmentation (Fig. 4g). The FN1218/6 deletion is allelic to the *d* allele in JI0073 and JI2202 (*P. abyssinicum*, a taxon that lacks axil ring pigmentation) (Supplementary Figs. 20-22), implicating these genes as corresponding to *D*.

The results presented here reveal the complexity of axil ring pigmentation regulated by *D*. There are multiple alleles of *D* within the *MYB* gene cluster, and many spontaneous conversions from one allelic form to another^50^, suggesting that it is the combination of alleles at several of these *MYB* genes which determines the presence, absence or pattern of this pigmentation. Both *a* and *a2* are epistatic to *d*, and we can postulate that the *MYBs* involved in the *D / d* phenotypes are part of a *MYB* (*D*) – *bHLH* (*A*) - *WD40* (*A2*) complex^12,68,69^.

#### A quantitative trait essential in pea breeding

Mendel examined the segregation of traits that have clear alternative states; he also noted that seed size (among other traits) differed between his parental lines, but considered that this quantitative difference was not suitable for his analyses. Seed size in pea defines some market classes such as the so-called ‘marrowfat’ types, with large irregular shaped seeds and a high protein content. Seed size has been the subject of QTL analyses^70–72^, and we have investigated this further within our diversity panel.

We discovered a significant novel locus on chromosome 2 that influences both pod width and hundred grain weight (Supplementary Fig. 23) and is in a similar location to a previously described seed size QTL in Medicago and pea^73,74^ (Fig. 4b, c). We designated this locus *Organ size 1*, *PsOs1*. Combining fine-mapping (Supplementary Fig. 24 and Supplementary Table 32) and differential gene expression analysis, we identified *Psat02G0011300* as a gene candidate for *PsOs1* (Supplementary Figs. 25-26), which encodes a SIAMESE-related protein (SIM/SMR), a cyclin-dependent protein kinase inhibitor (CKI), influencing cell division and enlargement during the cell cycle and consequently altering plant cell size^75^. Functional validation from the VIGS (Supplementary Fig. 27) approach, coupled with a transgenic overexpression line in Arabidopsis (Supplementary Figs. 28-29) demonstrate the key role of *PsOs1* in regulating seed weight and pod width.

## DISCUSSION

Despite the clarity of his 1866 paper, there is some dispute about what Mendel did. It has been argued that Mendel was not primarily interested in inheritance^76,77^, or that he had a pre-formed theory of inheritance that he sought to demonstrate, even to the extent of fabricating data to conform with his theory^78^. These views are mutually exclusive, and we reject them both^51,79^.

We have shown that variation in the genes underlying the seven pairs of contrasting traits that Mendel studied corresponds to a remarkable diversity of mutational mechanisms (Supplementary Table 33). There are several point mutations in *a*, one affecting the pattern of splicing and two different single nucleotide insertions affecting the reading frame, while *le* corresponds to an amino acid substitution caused by a missense mutation. The parchmentless mutation *p* corresponds to a single nucleotide substitution generating a premature stop codon, while insertion events of class I and class II transposons explain green cotyledons (*i*) and wrinkled seeds (*r*), respectively^21,23^. We have also uncovered additional novel types of variation, corresponding to DNA deletions that lead to loss-of-function, such as the remarkable case of *gp*, with a large DNA deletion upstream of *ChlG*, a promoter deletion in the *i-2* allele, the *fa* allele with a small deletion within an exon, and new alleles of *a* with one or more exons deleted. An unexpected discovery in this study was of the existence of an intragenic suppressor allele of *A* which implies that the *a* allele was in existence long enough for this unlikely second site mutation to have occurred. The earliest known mention of a white flowered pea in the 1300s^12^ most likely reflects the history of documentation rather than of this mutant allele. It is interesting that this intragenic suppressor mutation corresponds to a shift in the position of an intron, which is rarely identified, even in inter-specific comparisons of many genes^80^.

The biological processes these genes represent range from variation in the activity of enzymes in primary metabolism (*r, i, gp*), hormone interconversion (*le*), transcription factor regulation of secondary metabolism (*a*), to the regulation of cell fate during development (*p, fa*). It is noteworthy that the two green vs yellow phenotypic differences correspond to disruption of either the final step of chlorophyll synthesis (*gp*) or the first step of chlorophyll degradation (*i*) and this synthesis vs degradation difference accounts for which phenotype, green or yellow, corresponds to the dominant vs recessive allele. The elucidation of the biochemical and regulatory mechanisms underlying these genes are outside of the scope of this study, but the genomic and genetic discoveries and insights presented here are crucial to help us further understand Mendel’s pea traits. For example, based on the discovery of the fused aberrant transcripts arising from the *NLR-CHLG* genomic region, we propose that transcript stability is altered by transcriptional interference during chlorophyll synthesis or through a nonsense-mediated decay pathway, leading to an increased degradation rate of *CHLG* transcripts (Supplementary Notes). In addition, to confirm the gene identity of *V* and *Mfa*, more investigation in biochemical genetics is needed to elucidate the potential *Mfa-CIK-CLV3-WUS* regulatory network underlying the meristem defects.

A longstanding question in relation to Mendel’s pea work was whether the phenotypic variation he described corresponded to rare variants of genes which explain only a minor proportion of the genetic variation for that trait. Our GWAS analyses emphatically show that this is not the case, and indeed that in one case where genetic heterogeneity was expected (fasciation) the variation we detected corresponded to a single genetic locus (*Fa*), albeit with a previously unsuspected modifier locus (*Mfa*). There are three caveats to this claim. The first is that the parchmentless pod is (as has long been known) determined by either *P* or *V*, or the combination of these two distinct and independent genetic loci. A second caveat is that for green vs yellow cotyledon, there are clearly multiple GWAS peaks, albeit with lower significance than that of *I*. This probably reflects the influence of the seed maturation process on the penetrance of this phenotype (as was noted by Mendel in his 1866 paper). Finally, we observed an unusual feature of the GWAS peak corresponding to *Gp*, where there is a broad shoulder corresponding to most of the short arm of this chromosome. The reason for this is unknown.

This raises two general questions about GWAS analyses in defining genetic variation underlying traits. First, do broad GWAS peaks provide sufficient resolution to identify a manageable number of candidate genes? Second, how do the positions of significant GWAS signals correspond to previously described genetic variants? We have seen that for the seven Mendelian traits (and *D*), the GWAS peaks are significant, and all correspond well to the expected genetic loci. Furthermore, in our broad survey of many other agronomic traits for genotype-phenotype associations (Fig. 4a), nearly all the GWAS peaks correspond to the location of previously described genetic loci. This demonstrates that pea is an excellent model system for association genomics studies and GWAS is a suitable first step for trait-gene discovery and functional elucidation. The reliability of GWAS in pea is partly due to the fact that an unusually high proportion of pea genes are single copy^4^, and we established a high-quality genomic and phenotypic variation map from a global *Pisum* diversity panel, within which there is a rich reservoir of genetic diversity, as shown in this study. However, the pea genome is large and gene density is low throughout the chromosomes, maintaining a strong extended linkage disequilibrium. Presumably this is, in part, because of the strict inbreeding habit of pea.

We have shown how additional complementary approaches can narrow down these intervals to candidate genes. For the genes characterised, GWAS intervals alone were insufficient to delineate small sets of candidate genes. Additional resources such as specific biparental mapping populations, FN mutants, and functional validation are necessary. Future work requires innovative approaches and new technologies like the long-read DNA and RNA sequencing, a mature pea transformation system and targeted gene editing. These would help to examine in detail the multiple aberrant transcripts produced at the *gp* locus, transcriptional disruption by intronic LTR insertion in the *i-1* mutant, and genetic complexity of alleles at *D* due to gene redundancy within the *MYB* gene clusters, to further advance our understanding of Mendel’s traits.

The genomic, genetic and phenomic dataset from this large collection of *Pisum* accessions represents a permanent and invaluable resource. The very large numbers of genotype-phenotype associations we have found represent the beginning of a new phase of systematic trait dissection at the molecular and genetic level in pea. This study is essential for pea basic research, education in biology and genetics, and breeding practices.

## ACKNOWLEDGEMENTS

We thank Dale Sanders from JIC and Sanwen Huang from CAAS, who initiated this open and complementary collaboration; we thank Graham Moore for his invaluable support for this project. We thank Malcolm Bennett, University of Nottingham, UK and Jeffrey J. Doyle, Cornell University, USA, for their useful comments. We thank colleagues for assistance in pea field trial and phenotyping work from experimental stations across northern and southern China, Qiang Wang (HeiLongJiang (HarBin) Academy of Agricultural Sciences), and Yazhi Qin (Shenzhen) AGIS-CAAS Experimental Station. This work was supported by the Program for Guangdong “ZhuJiang” Introducing Innovative and Entrepreneurial Teams (2019ZT08N628), the National Natural Science Foundation of China (32022006), the Agricultural Science and Technology Innovation Program (CAAS-ASTIP-2021-AGIS-ZDRW202101), the Shenzhen Science and Technology Program (AGIS-ZDKY202002), the National Key Research and Development Program of China (2023YFF1000100), and the National Key R&D Program of China (grant number 2023YFA0914600) to S.C. We thank Emily Jones and Eilidh Crawford for plant phenotype data, and Martin Trick and Simon Griffiths for valuable discussions at the John Innes Centre (JIC). We also thank the John Innes Centre (JIC) NBI Computing Infrastructure for Science and JIC Bioinformatics groups for support in data handling and analysis, the JIC Field Trials and Horticultural Services teams for support with field and glasshouse experiments, the Molecular Genetics, Genotyping and DNA Extraction Platforms for support in experimental biology, and Bioimaging and Scientific Photography Platforms for phenotype visualisation. The work in the UK was possible due to the long-term investment of the UK Research Infrastructure Biotechnology and Biological Sciences Research Council (UKRI-BBSRC) through Institute Strategic Programme (ISP) grants, Institute Development Grant funds, and the Germplasm Resources National Capability Programme (BBS/E/J/000PR8000) and the National Bioscience Research Infrastructure grant (BBS/E/JI/23NB0001). We also acknowledge support from UKRI-BBSRC grants BB/J004561/1, BB/W510695/1 and BBS/E/J/000PR799, the UK Department for Environment, Food, and Rural Affairs (Defra) through the Pulse Crop Genetic Improvement Network (grants CH0103 and CH0111) and the Provision and Maintenance of the Pea Genebank to Facilitate R&D Need grant (C5515), and JIC through its Institute Strategic Fund. We acknowledge the source of the TILLING mutant used in the analysis of *gp* as UMR1403-INRAE-IPS2, UMR1347-Agroecologie, France and the EU Framework Programme 6 GRAIN LEGUMES.

## AUTHOR CONTRIBUTION

S.C. and N.C. conceived, designed, coordinated and managed the project; S.C., F.C., Y.S., and M.J. led the genomics, association genetics analysis, field trial and phenotyping, mapping genetics and functional validation work in China; C.F., Y.S., M.J., and H.C. led population genomics, bioinformatics pipeline development, haplotype-phenotype and association genetics analyses under S.C.’s supervision; B.C., Y.S., L.L., L.W. and Yu.S. led the field trial and phenotyping, gene cloning, RNA-seq, and VIGS gene silencing for *Gp*, *Fa*, *D* locus and *Os1*. S.C., F.C. and B.C. led the gene identity and variation discovery for *Gp*, *P*, *V*, *Fa*, as well as for the four cloned genes *R*, *I*, *A* and *Le*. Bo.S., H.Z., X.X.Z., X.L., and Z.H. participated in the whole-genome re-sequencing and bioinformatics analysis. X.W.D. participated in bioinformatics analysis and computation support from Peking University. J.H. and N.E. undertook genetic and genomic analyses of *A*, *D*, *Gp* and *Fa* including the allelism tests and TILLING validation of the nature of *gp*. M.D., A.F. and C.LeS. identified and provided *ChlG* TILLING mutant seeds. B.S. coordinated and managed bioinformatic and genomic data analysis at JIC. N.E. commented upon experimental results and assisted in data analysis of all the project results. N.C., C.D., N.E. and L.W. selected the germplasm panel. M.Vic. and R.W. assembled the JI0015 genome reference under B.S. supervision and together with J.C., C.LeS., A.B., M.D., C.S., C.M., R.S., M.Vig. and G.T. supported the *Gp* genetic and genomic explorations under the coordination of C.D. E.V., M.A., A.H., N.E., J.H., N.C., P.HP., and E.B. contributed to germplasm phenotyping the sequenced panel. M.A., L.S. and N.C. contributed through germplasm curation and R.H. by digitising germplasm data. S.C., N.C. and C.D. secured the project funding. S.C., N.E., F.C., J.H., M.J. and N.C. prepared the Figures, Extended Data Figures, Supplementary Figures and Supplementary Tables; S.C. and N.E. drafted and finalized the manuscript, with additional help from C.F., B.C., J.H., N.C. and B.S. All authors read and approved the final manuscript.

## COMPETING INTERESTS

The authors declare no competing interests.

## ADDITIONAL INFORMATION

Correspondence and requests for data and materials should be addressed to

## DATA AVAILABILITY

All whole-genome sequence data has been deposited at the National Genomics Data Center (NGDC) Genome Sequence Archive (GSA), with BioProject accession number PRJCA023166, and with SRA Accession ID: subCRA023387 (https://ngdc.cncb.ac.cn/gsa/, https://ngdc.cncb.ac.cn/gsa/s/49YdHBP5). Phenotyping data were given in Supplementary tables and long-term phenotype curation available on SeedStor (https://www.seedstor.ac.uk).

## GERMPLASM AVAILABILITY

All the germplasm described and used in this work is available to order from the John Innes Centre Germplasm Resources Unit (https://www.seedstor.ac.uk/). The 697 sequenced single seed derived JI *Pisum* Germplasm accessions are also available from the Agricultural Genomics Institute at Shenzhen, Chinese Academy of Agricultural Sciences.

## CODE AVAILABILITY

Code associated with this project is available at Github: https://github.com/ShifengCHENG-Laboratory/MendelPeaG2P

## ONLINE METHODS

### Plant Materials and Methods

#### Germplasm panel

A total of 697 accessions, maximising genetic diversity, were selected from the JI *Pisum* Germplasm Collection for this study. (Supplementary Table 1 and www.seedstor.ac.uk). These are also maintained at the Agricultural Genomics Institute at Shenzhen (AGIS), Chinese Academy of Agricultural Sciences (CAAS), China.

#### DNA extraction for whole-genome resequencing

Genomic DNA was extracted from approximately 50 mg leaf tissue of three-week old seedlings. Extraction used the oKtopureTM system (LGC Biosearch Technology) following tissue desiccation with silica for 48 h. A bespoke protocol was used with the following volumes per sample: 250 µl lysis buffer, 170 µl Binding buffer, 20 µl sbeadexTM suspension, 300 µl PN1 wash buffer, 300 µl PN2 wash buffer, 300 µl PN2 wash buffer (x3 wash cycles) and using 75 µl final Elution buffer. For each accession, a minimum of 6 μg of genomic DNA was used to construct a 150 bp paired-end sequencing library with an insert size of 500 bp following the manufacturer’s protocols (employing PCR-free methods), which was subsequently sequenced on the DNBSEQ Platform at BGI-Shenzhen resulting in ∼80 Gb clean reads with a coverage of ∼20X for each accession.

#### Phenotyping

DNA was extracted from a single plant whose seed was bulked up for progeny phenotyping. The diversity panel was planted in three different sites, Norwich, UK (52.62° N, 1.28° E), Shenzhen (Southern China, 22.61° N, 114.51° E) and Harbin (Northern China, 45.86° N, 126.83° E). In China, four rounds of phenotyping were conducted. Specific subsets of accessions and some F2 populations were grown indoors in the greenhouse of Shenzhen Agricultural Field Farm, with 16 hours of light/8 hours of darkness. Phenotypes collected at the three stations (2020 -2023), and a historical JIC phenotype dataset were curated in Seedstor (www.seedstor.ac.uk). In Shenzhen, peas were planted in winter (October) and harvested in March the following year, while in Norwich and Harbinthey were planted in spring (March to April) and harvested in August to October of the same year.

For the phenotyping of pod colour (green vs yellow podded lines), a field trial of three 1m^2^ microplots of 100 seeds each was sown in Spring 2023 where a 1:1 ratio of BC6 S3 *GpGp* and *gpgp* seeds (selfed seed of S2 homozygotes) were mixed and sown at random in each plot. At the pod filling stage, the *Gp* plants were tagged and at plot harvest seed was collected from individual plants to determine the the yield of *Gp* and *gp* homozygotes. Seeds were weighed and counted on a Data Count R25+ machine (data-technologies.com). Pod length and width were measured on 25 randomly selected pods. For the phenotyping of organ size, Pod width (PW) and hundred grain weight (HGW) were measured in mature pods of the F2 and F_2:3_ populations post-harvest. In the F2 populations, PW was assessed using 15 representative pods, divided into three groups of five, with the total width of each group measured sequentially. For the F_2:3_ populations, PW was determined using 5 representative pods, with their total width measured in a similar manner. HGW was calculated by randomly weighing 100 seeds from each accession, and repeating the process three times to obtain an average weight for each accession. Other more specific phenotypes were collected as described in Supplementary Table 27 and in line with published descriptors https://www.seedstor.ac.uk/search-phenotypes.php.

### Construction of the Pea Genomic Variation Map

#### Read Mapping, SNP calling and SNP annotation

The trimmed clean reads of each accession were aligned against the reference genome of pea (*P. sativum*) cultivar, ZW6^24,86^ and Caméor v1.0^4^, using BWA-MEM (v0.7.17) with default parameters^24,86^. Unmapped, non-unique and duplicated reads were filtered out using SAMtools^87^ (v1.9) and Picard (v2.20.3-SNAPSHOT) before variants were called by a standard pipeline of Genome Analysis Toolkit (GATK^88^, v4.1.2). SNPs were further filtered to remove low-quality variants defined as (1) SNPs with more than two alleles; (2) SNPs with QD<2.0, FS>60.0, MQ<40.0, SOR>3.0, MQRankSum<−12.5, ReadPosRankSum<−8.0; (3) SNPs with observed heterozygosity (H_obs_) exceeding the maximum calculated value (H_obs_max_) based on the Inbreeding Coefficient (F), where F was calculated as 1 - (H_obs_/H_exp_), with H_exp_ defined as 2p(1-p) using the frequency of the non-reference allele, and H_obs_max_ was determined as 10*(1-F_median_)*H_exp_ for variants with F>0 and MAF >0.05; (4) SNPs with missing rate >20% and MAF < 0.01. SnpEff^89^ (version 4.3t) was used to annotate the SNPs, and functional significance was then categorized based on their positions with respect to genes (intergenic regions, exons, introns, splicing sites, untranslated regions, upstream and downstream regions) and mutation consequences (missense, start codon gain or loss, stop codon gain or loss and splicing mutations).

#### Identification of Indels, gene PAV and gene CNVs, and SV

Small InDels (<=50bp) were called using GATK (v4.1.2) and filtered following the criteria: QD < 2.0 || low_QD || FS > 200.0 || high_FS || ReadPosRankSum < -20.0 || low_ReadPosRankSum before they were annotated using SnpEff (v4.3t). Read depth variation from read mapping analysis was used to identify gene presence and absence variation (PAV) and gene copy number variation (CNV) through normalization and correction in statistical analyses, following five steps: (1), mapped read depth at each gene was counted for each accession; (2), a correction for read depth variation (RDV) was applied, accounting for highly similar genes through all-vs-all CDS alignment using BLASTN. Recently duplicated genes were collapsed into representative genes to minimize depth bias, which were further normalized by dividing the corrected read depth of the gene by the average sequencing depth of the accession; (3) the distribution of read depth vs. GC content was used to correct read depth bias for each gene resulting from differential GC contents; (4), read depth variation was corrected for genomic regions with insertions or deletions in the genome reference; (5), subspecies-unique and shared CNVs were characterized by calculating the number of accessions with different copy numbers for each gene within each subspecies.

Different categories of structural variants (SVs: duplication, inversion, translocation, and large-scale deletion/insertion) were detected based on read mapping (read depth and read pair relationships) on PCR-duplicate-marked bam files using Delly^90^ (v 0.8.7) with default parameters; a summary of SVs identified is given in Supplementary Table 10.

#### Linkage disequilibrium (LD) analysis and Pea Haplotype map (HapMap)

A two-step LD pruning process was implemented to generate a high-quality core SNP dataset for the construction of a haplotype map^91^. Initially, SNPs were pruned based on linkage disequilibrium (LD) using PLINK^92^, with a window size of 10 kb, a window step of one SNP, and an r^2^ threshold of 0.8. A second round of LD pruning was conducted with a window size of 50 kb, a window step of one SNP, and the same r^2^ threshold of 0.8. For population LD-based haplotype analysis, the filtered SNPs were phased using Beagle (v 21Apr21.304)^93^. Subsequently, haplotype blocks were delineated utilizing PLINK with specific parameters (--blocks no-pheno-req --blocks-max-kb 1000 --geno 0.1 --blocks-min-maf 0.05). To merge adjacent blocks maintaining significant LD, D’ statistic values were calculated between all SNPs of consecutive blocks. If the lower quartile (Q1) exceeded 0.98, the adjacent blocks were merged. After filtering for the inbreeding coefficient, HAPPE^94^ was employed to identify haplotype clusters (haplogroups) for each block.

### Construction of Mapping Populations

#### JI0816 x JI2822 F2 population

Lines JI0816 and JI2822 (Supplementary Table 17), both of short stature, are maintained in the JI *Pisum* germplasm collection (https://www.seedstor.ac.uk/). JI0816, also known as WBH 1185, has pink flowers, a fasciated stem and yellow pods lacking pod parchment, corresponding to the mutant alleles *b, fa, gp* and *p*, respectively. JI2822, a recombinant inbred line derived from the cross JI0015 x JI0399, is wild type at these four loci. JI0015 and JI0816 share the *gp* allele, indicating that these two lines had a common parent, therefore segments of the genetic map are devoid of segregating alleles. 1000 F2 seeds from 9 F1 plants (JI2822 x JI0816) were sown at the JIC field station in Spring 2022. DNA preps from 942 plants were prepared from individual leaflets using the Qiagen DNeasy protocol (www.qiagen.com). Of these, 405 were genotyped using an axiom SNP array as described by Ellis et al^66^. The phenotypic and genotypic data are available in Supplementary Tables 17-19, and the sequences corresponding to the axiom markers are available in Supplementary Table 3 of Ellis et al^66^.

#### JI0015xJI0399 and JI2822xJI2233

Three populations have been used for mapping *Gp*. The first to be used was the previously recombinant described inbred population JI0015xJI0399 (Supplementary Table 20), later genotyped by Neogen UK, using an Infinium array as described previously^72^. The second was an F2 population derived from a cross between two of these RILs JI2822 *GpGp* and JI2833 *gpgp* which was screened using PCR for markers already mapped in JI0015xJI0399 in order to identify informative individuals (Supplementary Table 21). These, together with selected RILs with informative recombination events were genotyped with Axiom markers as described elsewhere (cite ref #90). *Gp* also segregates in the JI0816 x JI2822 F2 population as described above. The marker data are available in the supplementary file Gp mapping in JI0015 x JI0399 (Supplementary Table 20-21).

#### Other F2 mapping populations and Bulked Segregant Analysis (BSA)

We selected parental lines with contrasting pairs of traits to map genetic loci of interest in F2 populations using mapping by sequencing^95^ of bulked segregants. For genetic loci controlling uncharacterised Mendel traits; flower position (axial vs. terminal), pod colour (yellow vs. green), pod shape (inflated vs. constricted), crosses were made between Caméor (axial) × JI0814 (fasciated) and JI1995 (green pod) × JI2366 (yellow pod)F2 populations for the *P/V* loci (pod shape) were derived from four crosses, with JI0077 (*PPvv*), JI0466 (*ppVV*), JI0467 (*ppVV*) and JI0074 (*PPvv*) as male parents and JI1995 (*PPVV*) as the female parent. F2 populations for the *D* locus (one (*D^co^*) or two (*D^w^*) axial rings of anthocyanin pigmentation) were derived from three crosses, with JI0191 (*D^w^*), JI0794 (*D^w^*) and JI1669 (*D^w^*) as male parents and JI0328 (*D^co^*) as the female parent. F2 populations for the *Fn/Fna* loci (flower numbers) were derived from four crosses, with JI0441 (1fpn), JI2410 (3fpn), JI0745 (2fpn) and JI0746 (3fpn) as male parents and JI1995 (2fpn) as the female parent. The markers and BSA analysis of the F2 population is from^84^.

Approximately 300 plants from the F2 population of each of these crosses were planted in Shenzhen, ChinaWild type and mutant and bulked DNA samples were prepared by mixing equal amounts of DNA from 30 accessions with the dominant and recessive phenotypes, respectively. DNA was isolated from fresh leaves using the CTAB method^96^). 50X depth genome sequences for each of the parents and the bulked samples were generated. Short reads were aligned against the ZW6 reference genome using BWA-MEM (v0.7.17) and SNPs were identified using Samtools (v1.9). The variation dataset was analysed using the G’s value method of the QTLseqr package (v0.7.5.2).

#### Marker development and QTL mapping

The organ size-related quantitative trait locus (*PsOs1*) was fine-mapped using 21 Kompetitive Allele Specific PCR (KASP) markers for SNPs distinguishing accessions JI0074 and JI1995 after whole-genome resequencing in the candidate region. Each KASP marker was designed with two allele-specific forward primers (Supplementary Table 34) and one common reverse primer, based on 200 bp sequences upstream and downstream of target genic SNPs, following the standards of LGC Genomic Ltd., Hoddesdon, UK. The genetic linkage map was constructed using JoinMap V4.0 software. Windows QTL Cartographer V2.5 software facilitated inclusive composite interval mapping (ICIM) for identifying and analysing QTLs. A logarithm of odds (LOD) score of ≥3.0 was deemed indicative of a QTL.

### Genetic mapping of *Gp*

Green vs yellow pod colour segregates in the recombinant inbred (RIL) population derived from the cross between JI0015 (*gpgp*) and JI0399 (*GpGp*). The JI0015xJI0399 RIL population comprises 90 recombinant inbred lines, which, together with their parents were genotyped using an Infinium array (Neogen UK) that detected 13,204 biallelic SNPs. This enabled us to position 5,209 PsCam markers on a genetic map (JI0015xJI0399) and place *Gp* between the markers PsCam005046 and PsCam056084 (and their co-segregating markers). Additional mapping was undertaken, using an Axiom SNP array with 84,691 features^66^ of selected F2 progeny of a cross between JI2822 (*Gp*) and JI2833 (*gp*) together with RILs from JI0015xJI0399 crossings known to have recombination events at informative locations. JI2822 and JI2833 are both RILs from the JI0015xJI0399 population. With respect to the ZW6 assembly^24^, this placed *Gp* between the axiom markers AX-183865165 (Chr2:320968993) and AX-183571028 (Chr3:325580858) (JI0015xJI0399). Analysis of an F2 population derived from crosses between JI2822 (*Gp*) and JI0816 (*gp*) placed *Gp* between the axiom markers AX-183571050 (Chr3:321020350) and AX-183879077, (Chr3:324762848 see, Supplementary Table 17 JI0816xJI2822).

We performed different association genomics analysis for pod colours, including the SNP-based GWAS, LD-based haplotype GWAS, kmer-derived IBS-based haplotype GWAS, and the SV-based GWAS (Supplementary Fig. 6), all resulting in consistent and significant single GWAS peaks for pod colour located in the expected position of *Gp*, as seen in Manhattan plots (Supplementary Fig. 6).

#### Allelism tests for gp

Crosses were made between pairs of yellow-podded lines in the JIC germplasm collection (Supplementary Table 17). Seed and vegetative phenotypes were used to identify F1 progeny plants, and those accessions allelic, or non-allelic, to *gp* were identified by their yellow, or green pod colour, respectively.

#### Near isogenic lines for Gp vs gp

The JI0015 *gp* allele was introgressed into the Caméor background by sequential back-crossing and F1 progeny testing using a codominant PCR marker assay with one forward (25994_F) and two reverse (25994_15R and 25994_399R) primers (Supplementary Table 17). *Gp* (596 bp) and *gp* (688 bp) alleles were distinguished in a 35 cycle, 10s-30s-60s Touchdown PCR reaction that reduces the initial 62°C annealing temperature to 50°C in the first 10 cycles.

### Genome-wide Association Study

The multi-location and multi-season phenotypic dataset was used to perform genome-wide association studies with SNP matrix using GEMMA (v0.98.1)^97^, employing parameters (gemma-0.98.1-linux-static -miss 0.9 –gk -o kinship.txt and gemma-0.98.1-linux-static -miss 0.9 -lmm -k kinship.txt). The structural variation matrix was used to test for association with phenotypic variation for each of the selected traits using the same parameters as above. The haplotype map was used to test for association with phenotypic variation for each of the selected traits using RTM-GWAS^98^ with parameters (rtm-gwas-gsc –vcf in.vcf –out out.matrix and rtm-gwas-assoc –vcf in.vcf --covar out.matrix.evec --no-gxe).The results were visualized using in-house R scripts.

### Gene Functional Validation Experiments

#### Fast Neutron mutants

Several Fast Neutron mutants from a population described by Domoney et al. (2013)^99^, were included in this project. These were:

FN1453/1 *sil* - like

FN1091/4 lacking axil ring pigmentation, allelic to *d*

FN1218/6 lacking axil ring pigmentation, allelic to *d*

FN2026/7 *coch2* candidate

FN2073/5 lacking axil ring pigmentation, not allelic to *d*

FN2076/5 VicA FN deletion line

Crosses were made between pairs of lines lacking axil ring pigmentation (Supplementary Fig. 20) to test for complementation. Where possible, vegetative phenotypes were used to identify F1 progeny plants, and those accessions allelic, or non-allelic, to *d* were identified by the absence, or presence of pigmented axil rings, respectively.

#### Gene Silencing by Virus-Induced Gene Silencing (VIGS) assay

VIGS in peas was conducted in accordance with published methodology as described^100^. Primers specific to the VIGS-*PsOs1* constructs are provided in Supplementary Table 34. Spe I and EcoRI were used to linearize the pCAPE2 vector, which was kindly provided by Li et al. (2019)^101^, and corresponding fragments of targets were ligated into the vector to construct the vectors for VIGS assay. The negative control vector, pCAPE2-Con, was constructed in the same way by replacing the *PsCHLG* fragment in pCAPE2-*PsCHLG* with a 529 bp insert derived from a cDNA fragment of Bean yellow mosaic virus (GenBank accession no. AJ622899). The positive control vector, pCAPE2-PDS, targeting the phytoene desaturase gene, was also provided by Li et al (2019)^101^. These vectors were transferred into *Agrobacterium tumefaciens* (GV3101) and VIGS assays carried out following the protocol described by Constantin et al. (2004)^102^. Briefly, Agrobacterium strains carrying these vectors were shaken separately until OD600=1.2, followed by the collection and resuspension of the bacteria in injection buffer (NaCl: 10 mM/L, CaCl_2_: 10 mM/L, Acetosyringone: 0.1 mM/L) to a concentration of OD600=1.2. After resting for 2-3 hours, the solution of PCAPE2-target gene, PCAPE2-PDS (positive control), and PCAPE2-Con (negative control) was mixed with PCAPE1, separately, in equal proportions, and injected into 10-day-old compound leaves of the acceptant lines (Yunnan2070 or JI1995). After 24 h of darkness, they were transferred to long day conditions. New leaves of positive control plants bleached in about 10 days, indicating successful silencing of PDS. VIGS was employed for *PsCHLG*, *PsMYB16* gene within the *D* locus, and *PsOs1* which is described in detail below. The *PsCHLG*-VIGS fragment is, AATATATGGAAGATTCGTCTTCAACTTACAAAGCCTGTAACTTGGCCTCCATTAGTTTGGGGTGTAGTTTGTGGTGCTGCTGCTTCTG. Other gene-specific primers used for VIGS constructs are listed in Supplementary Table 34.

#### Transformation, gene overexpression and silencing of PsOs1

The *PsOs1* coding sequence of JI0074 was amplified (primers listed in Supplementary Table 34) and integrated into the pCAMBIA1305 vector, resulting in the pCAMBIA1305-*PsOs1*_JI0074_ construct. The plasmid was then introduced into *Agrobacterium tumefaciens* GV3101, which was subsequently employed to transform *Arabidopsis thaliana* (Col-0) via the floral dip technique. T_3_ generation homozygous transgenic *Arabidopsis* lines were selected for measurement of thousand-seed weight and the dimensions of elongated siliques.

#### GUS staining, GFP fluorescence observations and Flow cytometry

The pCAMBIA1305-*PsOs1*_JI1995_ vector was constructed using the same methodology, with primers detailed in Supplementary Table 34. Both vectors, pCAMBIA1305-*PsOs1*_JI1995_ and pCAMBIA1305-*PsOs1*_JI0074_, were introduced into the *Agrobacterium tumefaciens* strain GV3101. In these experiments, H2B-mCherry served as a nucleus marker. The agrobacteria were resuspended and infiltrated into *Nicotiana benthamiana* leaf epidermal cells using an infiltration buffer consisting of 10 mM MES (pH 5.6), 10 mM MgCl_2_, and 150 μM acetosyringone, at an OD_600_ of 0.8. Fluorescence was observed 48 hours after infiltration using a confocal laser-scanning microscope.

To compare the promoter activities of JI0074 and JI1995, we cloned sequences 3000 bp upstream of the coding region and inserted them into pCAMBIA1300-GUS, resulting in the constructs Pro_JI0074_-GUS and Pro_JI1995_-GUS. These were expressed in tobacco leaves and subsequently stained using a GUS Staining Kit (Coolaber Biotech, Beijing, China). GUS activity was quantified using the GUS Gene Quantitative Detection Kit (Coolaber Biotech, Beijing, China). For a detailed examination of *PsOs1* expression patterns in *Arabidopsis*, various *Arabidopsis* tissues were sampled from Pro_JI0074_-GUS transgenic plants. Post-ethanol decolorization, observations and photographs were taken under a microscope. Details of the primers used are provided in Supplementary Table 34.

Intact nuclei from pea pods were isolated using LB01 lysis buffer (Coolaber Biotech, Beijing, China), followed by RNA removal and subsequent PI staining. The nuclei were then quantified using a CytoFLEX flow cytometer. A minimum of 20,000 nuclei were counted for each sample, and each experiment was replicated at least three times. Data analysis was conducted using FLOWJO software, and representative images were presented. The endoreduplication index (EI) was calculated using the formula: EI=[(0×% 2C)+(1×% 4C)+(2×% 8C)+(3×% 16C)+(4×% 32C)]/100.

#### Anatomical studies and transmission electron microscopic (TEM) observation

Upon sampling, the shoot apices of Caméor and *fa* mutant line JI0814, and the pod walls of JI0074 and JI1995 were immediately preserved in FAA fixative. Paraffin sectioning was performed following established methodologies. Staining was conducted using safranin and fast green (JI0074 and JI1995) and Toluidine blue (Caméor and JI0814). Prepared slides were scanned using a NanoZoomer, and cell quantification was carried out using NDP.view2 software.

For TEM studies, pea leaflets and pods (18 days after flowering) were removed from BC3 S2 *gpgp* and *GpGp* plants, after 9 h of daylight. Tissue (1mm^2^) pieces were placed in a solution of 2.5% (v/v) glutaraldehyde in 0.05M sodium cacodylate, pH 7.3 for fixation. Samples were left overnight at room temperature, then processed for embedding (Leica EM TP embedding machine Leica, Milton Keynes, UK) by washing out the fixative with three successive 15 minute washes in 0.05M sodium cacodylate, followed by fixation in 1% (w/v) OsO_4_ in 0.05M sodium cacodylate for 2 h at room temperature. After three, 15 minute washes in distilled water, samples were dehydrated in an ethanol series (30%, 50%, 70%, 95% and two changes of 100% ethanol), then infiltrated with LR White resin (London Resin Company, Reading, UK) by successive changes of resin:ethanol mixes at room temperature (1:1 for 1 h, 2:1 for 1 h, 3:1 for 1 h, 100% resin for 1 h, then 100% resin for 16 h, and 100% resin for a further 8 h). Samples were polymerised in LR White resin at 60°C for 16 h, then sectioned with a diamond knife (Leica UC7 ultramicrotome, Leica, Milton Keynes, UK). Ultrathin sections (approximately 90nm) were placed on 200 mesh formvar and carbon-coated copper grids (Agar Scientific, Stansted, UK). Sections were stained with 2% (w/v) uranyl acetate for 1 h and 1% (w/v) lead citrate for 1 minute, washed in distilled water and air dried. Grids were viewed in a FEI Talos 200C transmission electron microscope (FEI UK Ltd, Cambridge, UK) at 200kV and imaged using a Gatan OneView 4K x 4K digital camera (Gatan, Cambridge, UK) to record DM4 files.

### RNA-seq and Gene Expression

#### RNA extraction and Pea transcriptome

At China, plant tissues (seed, root, nodule, leaflet, stem, flower, pod, stipule, tendril and apical bud) at different development stages (seedling, flowering and podding) were collected and fixed in Trizol before RNA extraction. Tissues were ground in liquid nitrogen and the FastPure Universal Plant Total RNA Isolation Kit (Vazyme, Nanjing, China) was used to extract total RNA, the quality of which was assessed by gel electrophoresis. For each sample, we performed short read RNA-sequencing using the DNBSEQ Platform at BGI group Shenzhen to generate 6-8 Gb raw RNA reads for each accession.

At JIC, RNA was prepared from young developing pods (flat pod stage, ∼60-70 mm in length) of each of the parental and RI lines derived from the cross between JI0015 (*gpgp*) and JI0399 (*GpGp*). Developing seeds were removed from the pods which were then rapidly frozen in liquid nitrogen. High-quality RNA lacking genomic DNA was extracted from 97 individual pod samples, using a Spectrum™ Plant Total RNA Kit (Sigma-Aldrich), and used for RT-PCR and RNA-seq experiments focussed on the identification and characterisation of gene candidates for *gp*. For the latter analysis, green-podded and yellow-podded RILs (95 in total) were assigned to three groups for each phenotype, ensuring that lines with contrasting plant phenotypes (e.g. plant height) were randomly distributed among the replicate groups (G1, G2 and G3 for green-podded RILs; Y1, Y2 and Y3 for yellow-podded RILs, with 15-17 RILs per pool). Equal amounts of RNA from every line within a group were pooled. RNA-seq (Illumina HiSeq4000) and initial bioinformatic analyses were carried out by the Earlham Institute, Norwich, UK.

#### Quantitative real-time PCR (qRT-PCR)

Total RNA was reverse transcribed to cDNA using Vazyme’s HiScript III First Strand cDNA Synthesis Kit (+gDNA wiper). RT-qPCR analysis was conducted using Vazyme’s Taq Pro Universal SYBR qPCR Master Mix, employing specific primers, with *PsACTIN* serving as the internal standard. Expression levels of genes were quantified relative to the control based using 2^-ΔΔCT^ method. Results represent the mean ± SD from three separate biological experiments. The primers used for RT-qPCR primers used are provided in Supplementary Table 34.

### Statistical Methods

#### General Statistical Analysis

Statistical analyses were conducted in R software suite (version 4.2, https://www.r-project.org/) unless otherwise stated. Two-tailed Students’ t-tests in the analyses of the phenotypes, such as seed weight and pod width, between different accessions were performed using the ‘t.test’ package in R software (v4.2). The correlation between different traits were tested by calculating the coefficients of Pearson correlation, as well as the P values, using the ‘cor.test’ package, with the method set to “Pearson’ for the correlation analyses between quantitative traits. Traits collected at different locations and in different years were analysed by calculating their rank correlations by setting the option ‘method’ to ‘Spearman’. The correlation between qualitative traits was assessed using the chi-square test using the ‘chisq.test’ package in R. Gene expression levels in different lines or tissues under different treatments was analysed using DESeq2^103^, in which the genes with a false discovery rate (Bonferroni) lower than 0.01 were defined as significantly regulated genes.

Principal component analysis (main text Fig. 1) was performed on the PLINK distance matrix using an Excel add-in downloaded from RIKEN, now available at https://systemsomicslab.github.io/compms/others/main.html#Statistics.

#### Population Structure Analysis

The core high-quality SNP dataset was used for population structural analyses. PCA and t-SNE analyses were first performed using beta Python modules sklearn.decomposition and sklearn.manifold. ADMIXTURE^104^ (version 1.3.0) was employed to analyse the population structure, with K increasing from 2 to 16.

Genetic differentiation (*F*st) and nucleotide diversity (π) were calculated with VCFtools (version 0.1.13). *F*st scores were calculated within a nonoverlapping 100-kb windows and π was calculated for each individual site and averaged across the genome for each group. LD was calculated on SNP pairs within a 500-kb window using PopLDdecay^105^(version 3.31; https://github.com/BGI-shenzhen/PopLDdecay) and the decay was measured by the distance at which the Pearson’s correlation efficient (r2) dropped to half of the maximum. Splits Tree analysis of the PLINK distance matrix was performed using SplitsTree4^81^.

---

**Extended Data Fig. 1.**
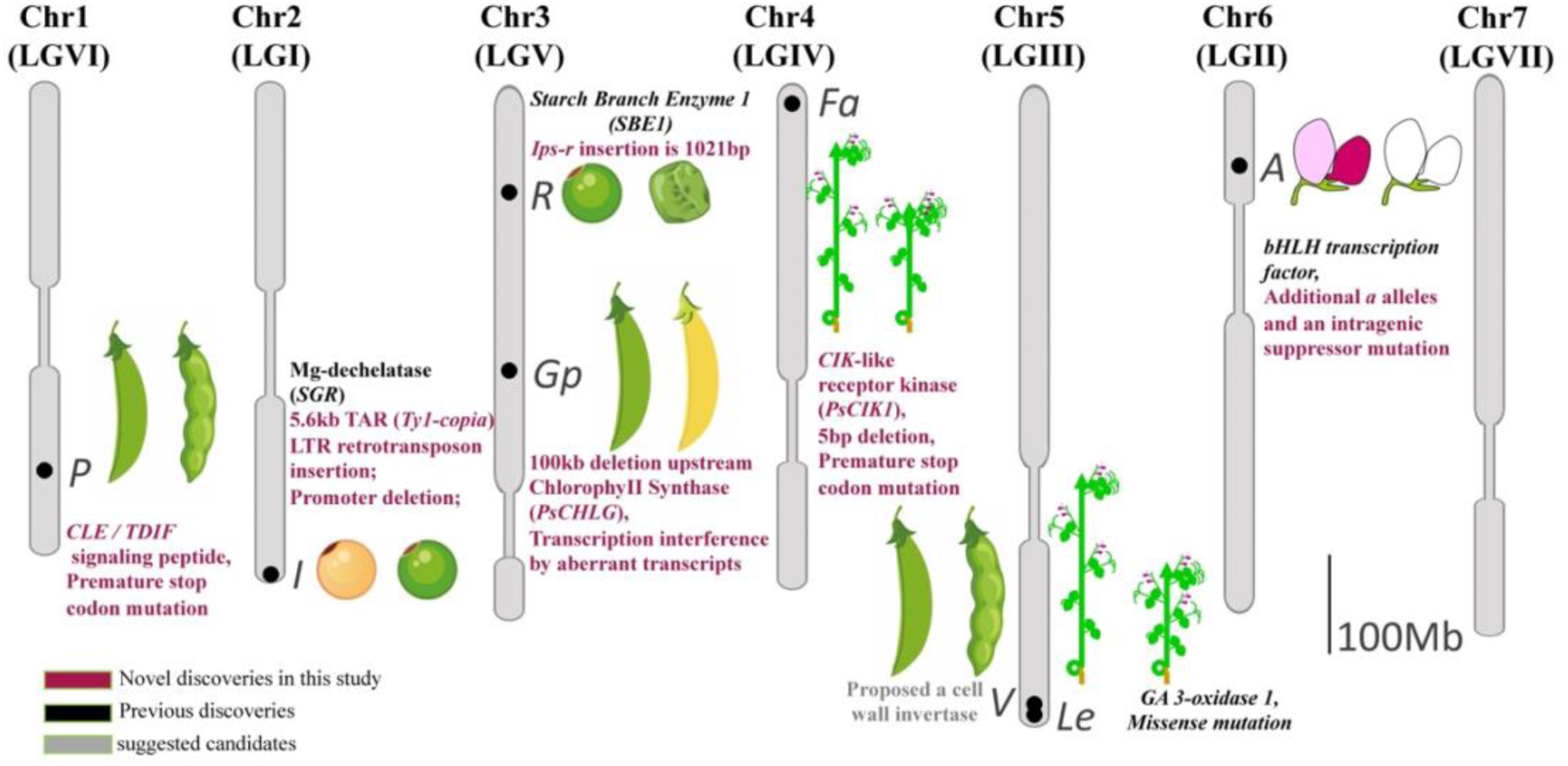
A schematic illustration of the genetic loci for each of Mendel’s seven traits plotted along the seven chromosomes (linkage groups). The previously cloned genes (*R*, *I*, *A*, *Le*) are annotated in black text, while the three remaining genes with gene identity and variations elucidated in this study (*P*, *Gp*, *Fa*) are highlighted in red text. The proposed gene candidate for *V* is highlighted in grey as this awaits more experimental data analysis. Difference in the form of the ripe pods on chromosome 1 (LGVI, *PP*/*pp*) and 5 (LGIII, *VV*/*vv*); Yellow versus green cotyledons (*II*/*ii*) on chromosome 2 (LGI); round seed versus wrinkled seed (*RR*/*rr*) and the colour of unripe pod (*GpGp*/*gpgp*) on chromosome 3 (LGV); difference in the position of the flower (*FaFa*/*fafa*) on chromosome 4 (LGIV); tall versus dwarf plants (*LeLe*/*lele*) on chromosome 5 (LGIII); seed coat (and flower) colour (*AA*/*aa*) on chromosome 6 (LGII).

**Extended Data Fig. 2.**
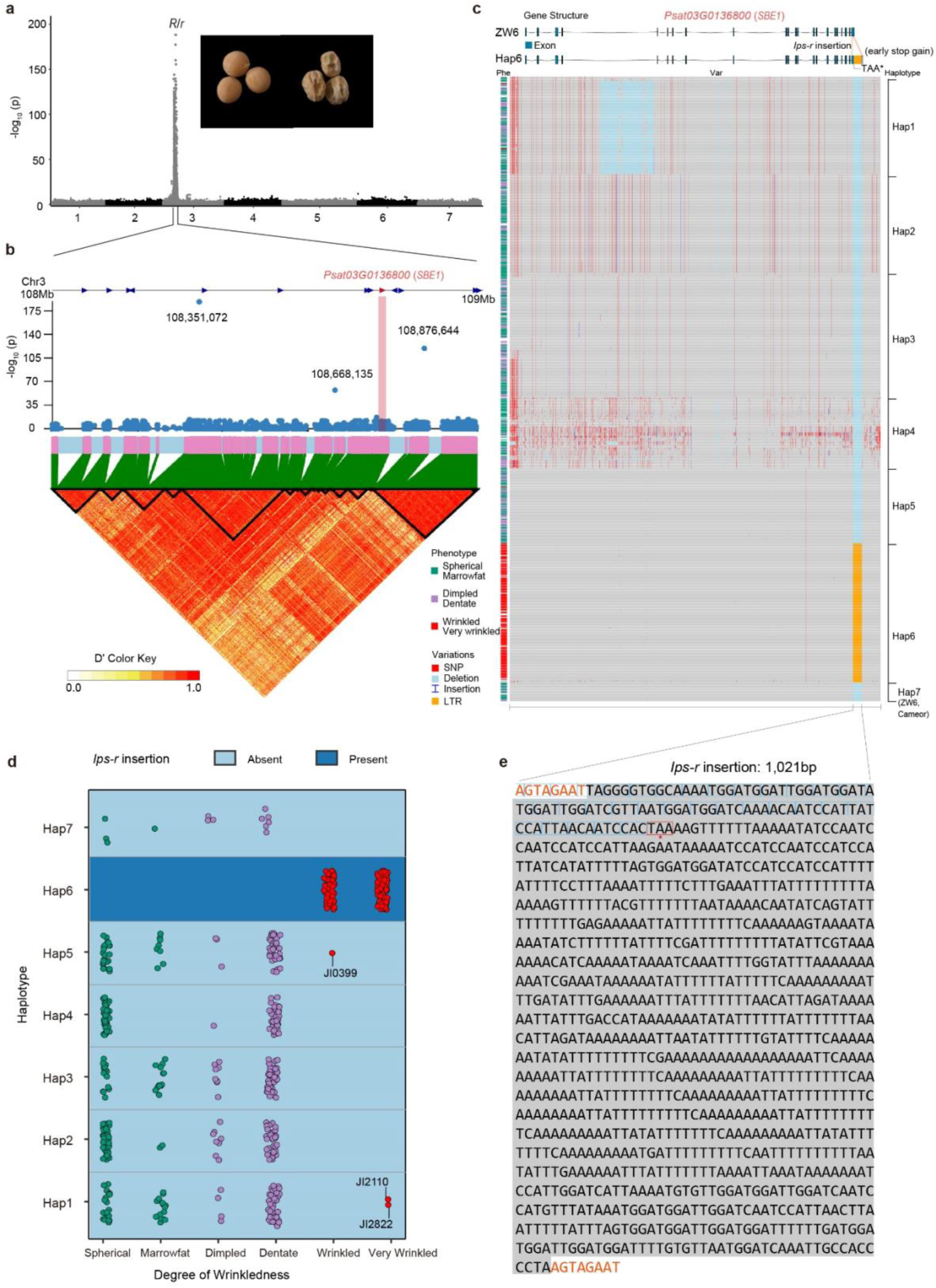
Haplotype-Phenotype association study for seed shape (round vs. wrinkled). **a,** Manhattan plot of GWAS based on the ZW6 genome reference explaining the round vs wrinkled phenotype, as illustrated. The single strong signal is consistent with the *R* locus located on the long arm of Chr3. **b**, Local detail of the genomic interval shows the gene list, with the *Starch Branching Enzyme 1* gene (*SBE1*, *Psat03G0136800*, chr3:108,732,329-108,770,718^23^) highlighted in red. The local linkage disequilibrium map is plotted below where the data points corresponding to *SBE1* (it is at 108,732,329-108,770,718) are marked in red; the significance values of these data points are quite low indicating that the SNP variants here are not causative and presumably do not distinguish the *r* alleles from the wild type *R* progenitor. **c**, a population-based haplotype clustering and haplotype-phenotype association analysis of *SBE1*, showing that most of the accessions with wrinkled seeds are clustered in Haplotype 6 which is consistent with the causal variation of *Ips-r* insertion event (marked in orange colour) in the last exon. Note that this haplotype does not have unique SNPs. A few accessions with wrinkled seeds distributed elsewhere are caused by the *rb* allele^29,82^ (JI0399, JI2822) or the previously described variant in JI2110, cv Kebby (see also panel d). **d**, the distribution of the different phenotypes (which were classified into six categories: Spherical (green dots), Marrowfat (green dots), Dimpled (purple dots), Dentate (purple dots), Wrinkled (red dots), and Very Wrinkled (red dots)) recorded in this study, corresponding to the different haplotypes given in panel c. **e**, the full sequence of the *Ips-r* element (1,021 bp), the 8bp site duplication (AGTAGAAT) sequences bounding the insertion event are highlighted in red. The extension of exon 22 into *Ips-r* is indicated by the blue boxes around the codons and the premature termination codon is boxed in red.

**Extended Data Fig. 3.**
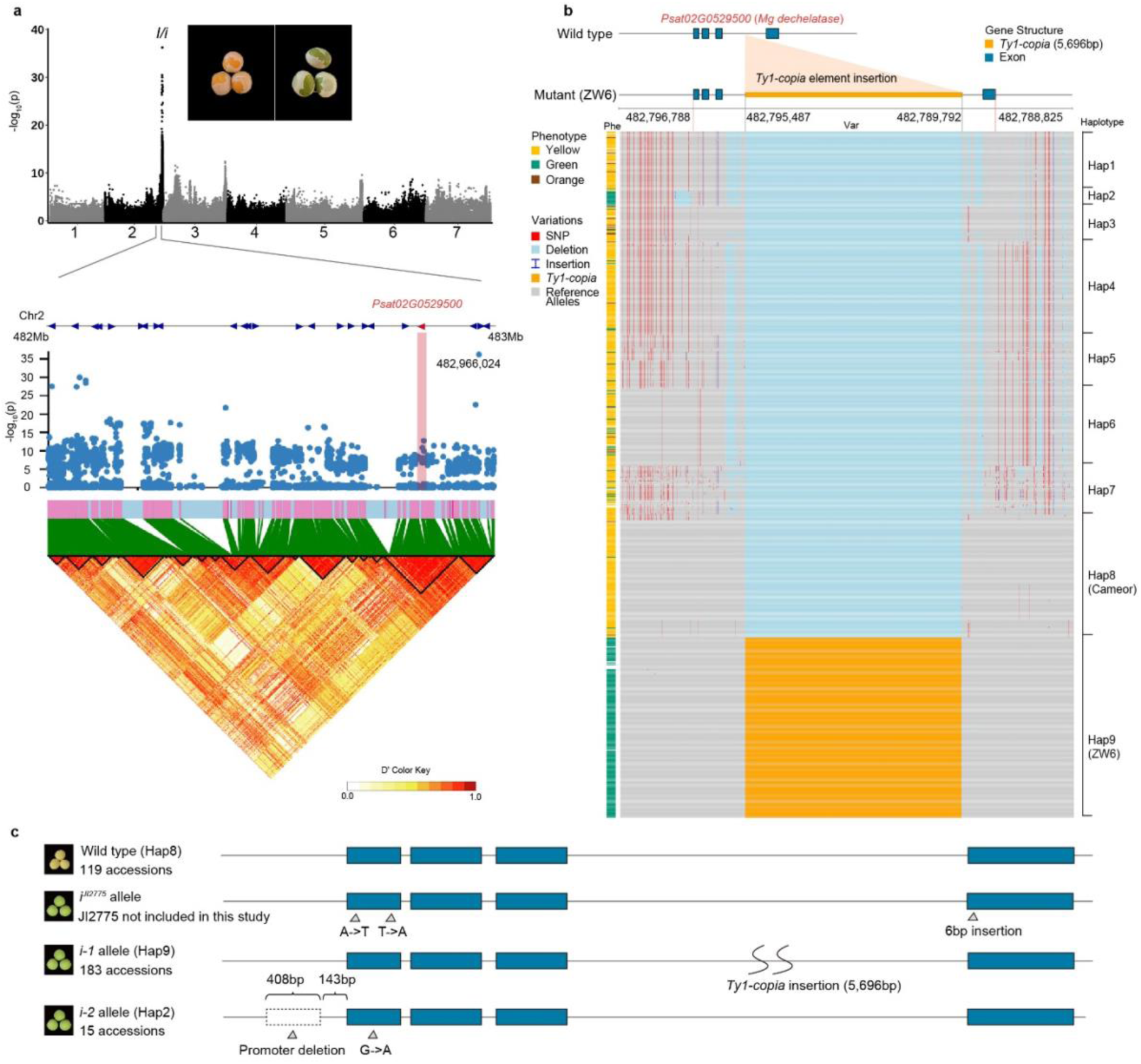
Haplotype-Phenotype association study for cotyledon colour (yellow vs. green). **a,** Manhattan plot of GWAS based on the ZW6 genome reference, where the strongest signal is consistent with the *I* locus located on the short arm of Chr2. The genomic interval is a narrowed down into a 1Mb region around the *Stay Green* gene (*SGR*, *Psat02G0529500*) highlighted in red. The local linkage disequilibrium map is shown below. The significance value of the data point from the local GWAS corresponding to *I* is low; the causative variation is an indel and the SNPs do not distinguish the *i* allele from its *I* progenitor. **b**, a population-based haplotype clustering and haplotype-phenotype association analysis of *SGR*, showing that most of the accessions with green seeds are clustered in Haplotype 9 which corresponds to the *Ty1-copia* insertion event in the last intron between exon4 and exon5. Note that this haplotype does not have unique SNPs. Haplotype 2 corresponds to a promoter deletion event that presumably disrupts the expression of *SGR*. Some of the other accessions with green seeds distributed elsewhere are possibly caused by other genes or by premature maturation of the seeds. The *Ty1-copia* insertion is marked in yellow both within the gene structure (top) and in the haplotype heatmap (in the middle of Hap9). **c**, *SGR* gene structures identified in this study and previously^21^. In this study only two allelic forms of *i* were found; *i-1* in Haplotype 9 and *i-2* in Haplotype 2.

**Extended Data Fig. 4.**
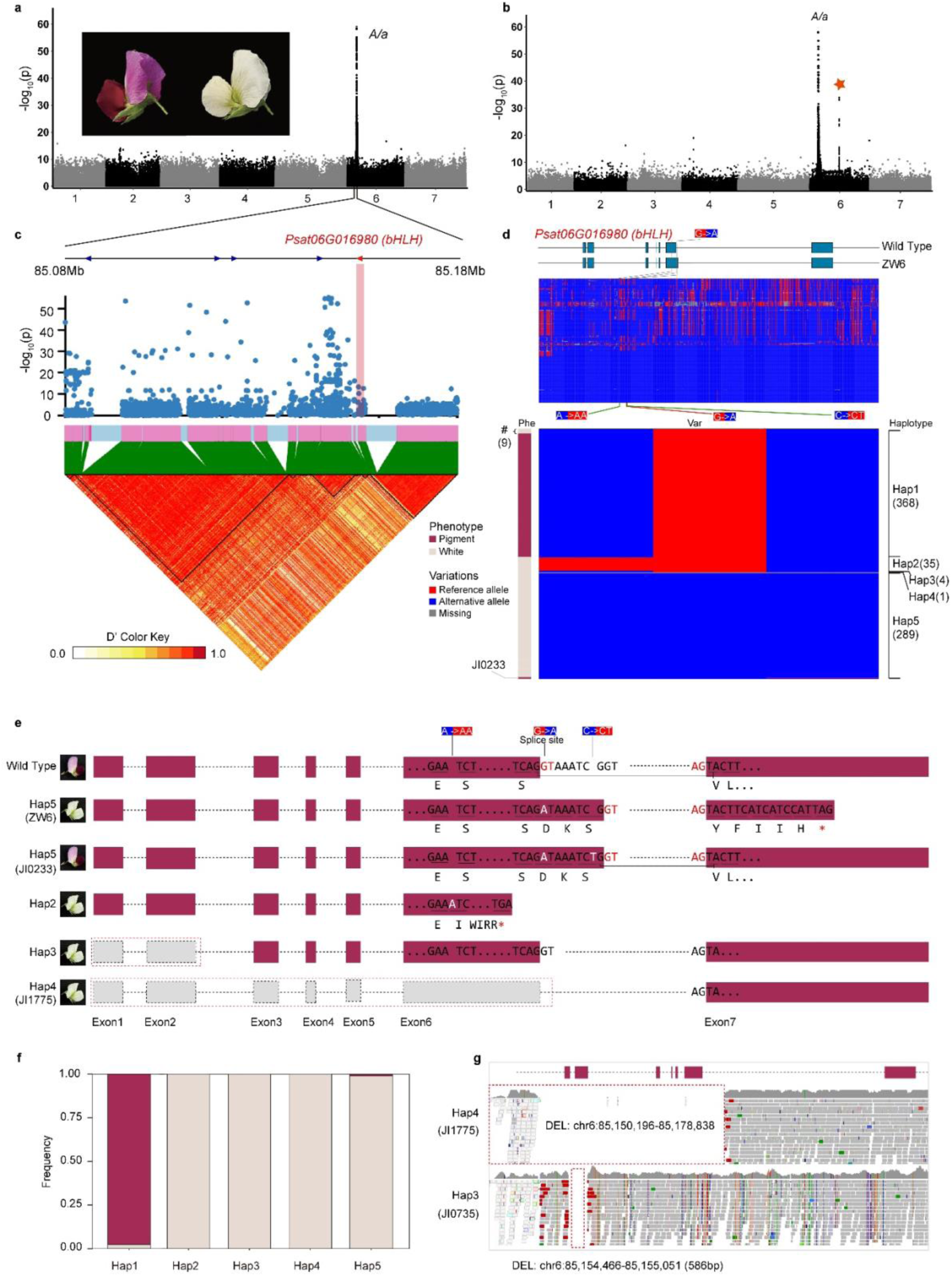
Haplotype-Phenotype association study for flower colour (pigmented vs. white). **a**, Manhattan plot of GWAS with respect to the ZW6 genome reference, showing a single strong signal which is consistent with the *A* locus located on the short arm of Chr6. **b**, Manhattan plot of GWAS with respect to the Caméor v1a genome reference, showing a second signal (marked with a red star) found in the middle of Chr6 which, with reference to the genetic map^66^, was attributed to an assembly error. **c**, Local detail of the genomic interval showing the gene list with the basic Helix-Loop-Helix (*bHLH* transcription factor, *Psat06G0169800*) highlighted in red. The local linkage disequilibrium map is shown below. **d**, a population-based haplotype clustering analysis of the *bHLH* gene. Upper panel for the whole gene, lower for panel functionally relevant SNPs. Five different haplotypes are indicated, most of the purple flowered lines (as shown in the left bar) belong to Hap1 carrying the G of the wild type intron 6 splice donor site^12^. Haplotypes 2-4 are mutant types with white flowers: The Hap2 allele has an additional A in exon 5, as originally described for JI1987^12^; Hap3 corresponds to the deletion of exons 1 and 2. Hap4 has a deletion of exons 1 to 6. Hap5 (including the ZW6 reference genome) is the most common mutant type with white flowers and carries the G to A substitution at the intron 6 splice donor site first described in Caméor^12^. However, within Hap5, there is one exception (JI0233) which carries this G to A substitution but has fully pigmented flowers (see main text). It is worth mentioning that the coding sequence (the *bHLH* transcript) is on the – strand in the reference genome (ZW6), here we standardize the comparison between the wild-type and the mutant types. **e**, Gene structures corresponding to the variants described in panel d. **f**, frequency distribution of the phenotypes (pigmented/purple vs. white) for each of the haplotypes. **g**, read mapping to confirm the exon deletion events that disrupt gene function conferring white flowers of Hap3 and Hap4 as shown in panel d/e.

**Extended Data Fig. 5.**
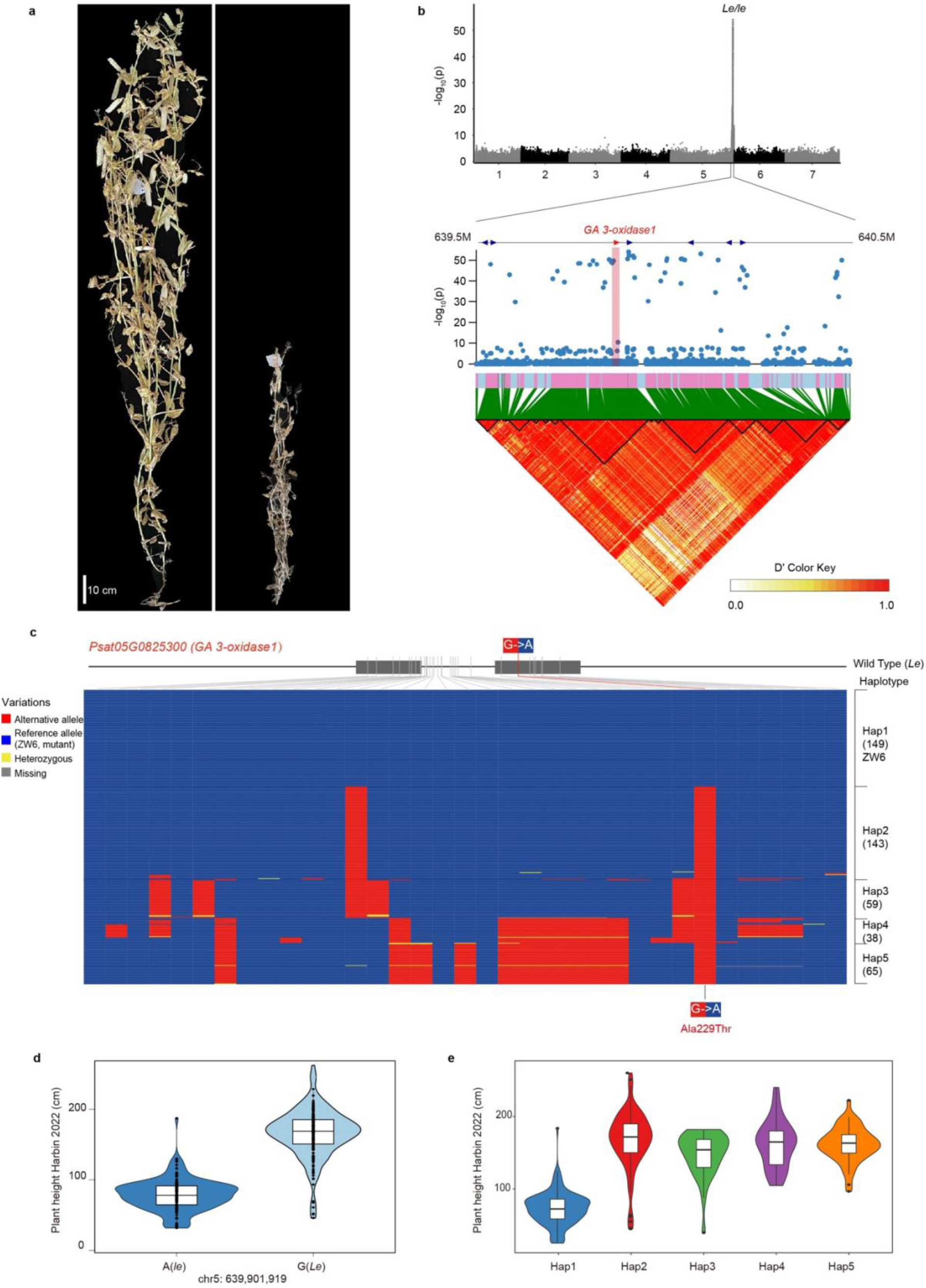
Haplotype-Phenotype association study for plant height (tall vs. dwarf). **a,** Two images showing the contrasting traits of stem length (long vs. short). **b**, Manhattan plot of GWAS, showing a single strong signal consistent with the *Le* locus located on the short arm of Chr5, based on the ZW6 genome reference. Local detail of a 0.5Mb region within this genomic interval including the *GA 3-oxidase1* gene^18,19^, *Psat05G0825300*, highlighted in red. The local linkage disequilibrium map is shown below. **c**, a population-based haplotype clustering analysis of the *GA 3-oxidase1* gene, showing five different haplotypes. Most accessions with short stem length are clustered into Hap1 carrying the previously described G to A mutation^18,19^, all the other haplotypes (Hap2-5) have the G nucleotide. The reference genome (ZW6) belongs to mutant type in the Ala229Thr substitution (nucleotide G to A) position. **d**, distribution of the phenotypes (plant height, Harbin location) corresponding to accessions carrying the mutant (A, *le*) or the wild type (G, *Le*) allele; **e**, distribution of the phenotypes (plant height, Harbin location) corresponding to the different haplotypes (Hap1-5).

**Extended Data Fig. 6.**
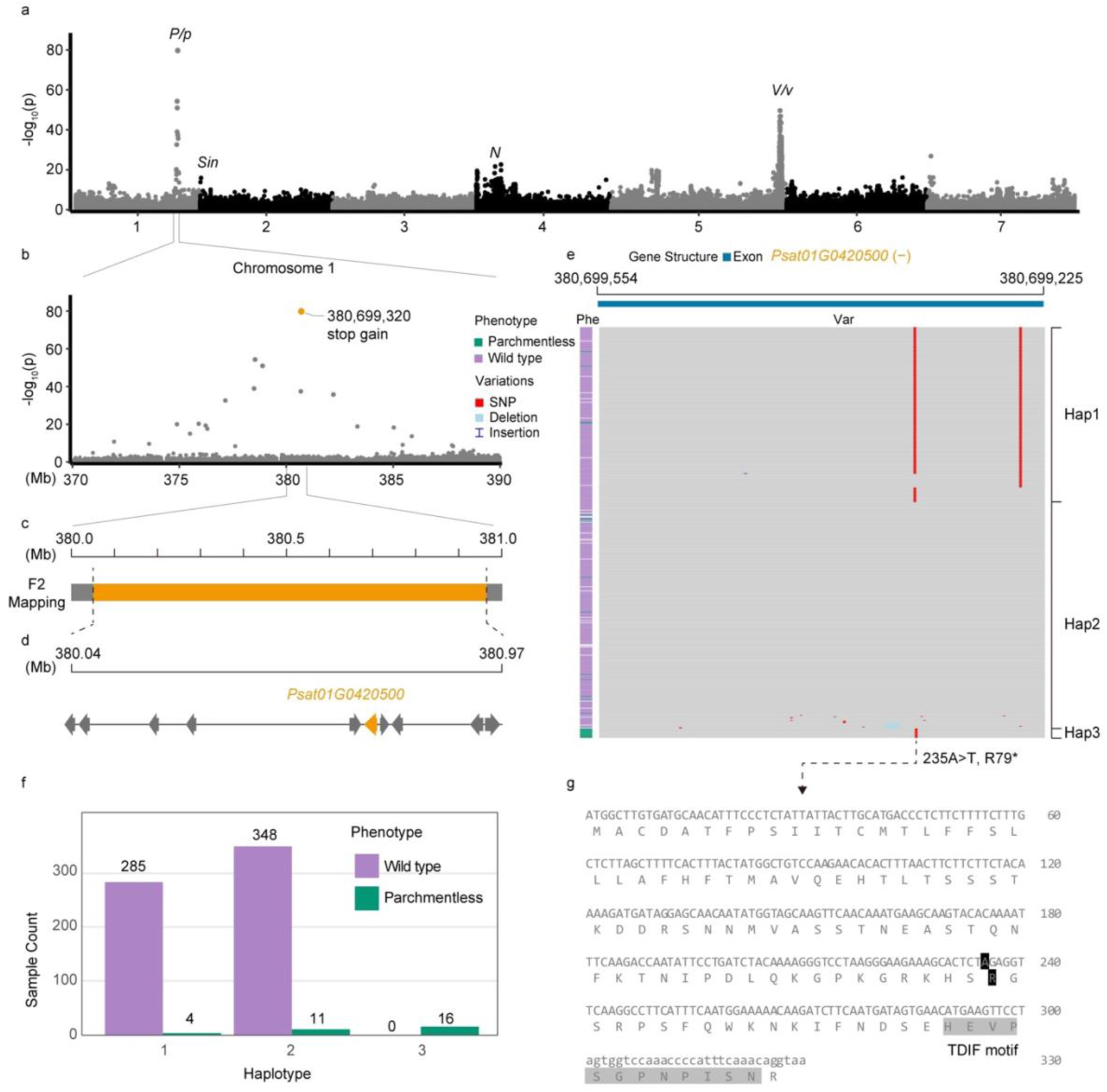
Gene identity and causal variations underlying parchmentless pods (*P* vs. *p*). **a,** Manhattan plot of GWAS based on the ZW6 genome reference for parchmentless pods (See also Fig. 2b of the main text). **b,** Close-up Manhattan plot of the most significant region in panel a. **c,** F2 mapping genetic interval from the cross JI0816xJI2822, showing the mapped locus between the markers AX-183563747 and AX-183563750 (chr1: 380049894-380967975) (Supplementary Table 17-19). **d,** Map of gene positions within the *P* interval with *Psat01G0420500* encoding a tracheary element differentiation inhibition factor CLE41/44 indicated in yellow. **e,** Allelic/haplotype variation for *Psat01G0420500*. Note that Hap1 has a silent A to C transversion at chr1_380699321, close to chr1_380699320 of Hap3, where the T to A transversion is responsible for the Arg79* nonsense mutation. **f,** Haplotypes of *Psat01G0420500* corresponding to accessions with ‘parchmentless’ phenotypes. **g,** Predicted amino acid sequence of *Psat01G0420500* indicating the position of the Arg79* mutation in relation to the TDIF^83^ motif.

**Extended Data Fig. 7.**
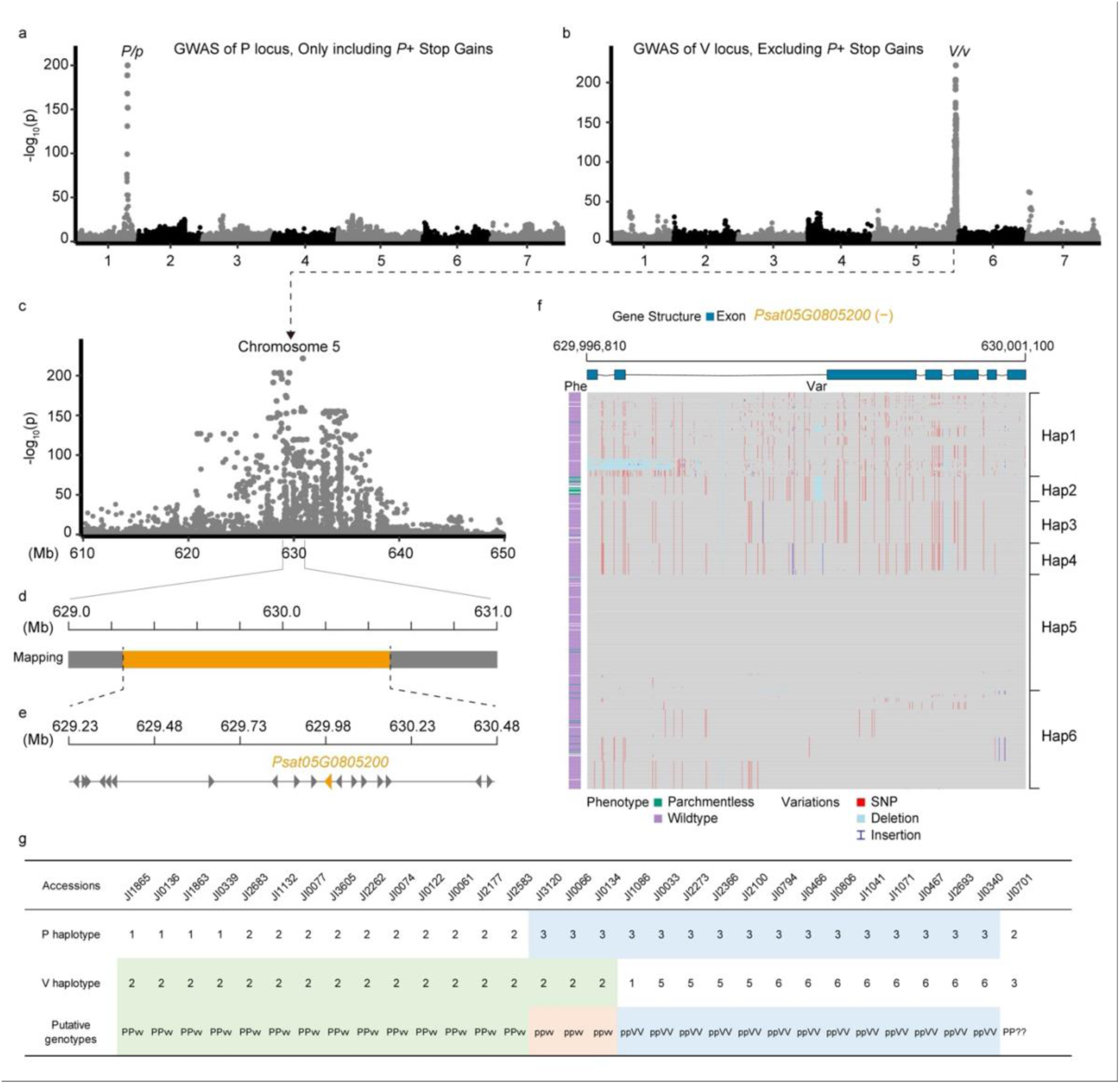
*V* and parchmentless pods. **a**, Manhattan plot of GWAS based on the ZW6 genome reference from a subset of accessions that include only the lines carrying the R79* allele (haplotype 3 in Extended Data Fig. 6 panel e) of gene *Psat01G0420500* and wild type accessions (i.e. no *vv* mutants), showing the *P* GWAS signal but not the *V* GWAS signal. **b**, GWAS from a subset of accessions (contrary to panel a) that exclude the lines carrying haplotype 3 (Extended Data Fig. 6 panel e) of gene *Psat01G0420500*, (i.e. no *pp* mutants); therefore, no *pp* mutants but only the *vv* mutant, showing only the *V* GWAS signal but not the *P* GWAS signal. **c**, Close-up of the local details of the chromosome 5 GWAS peak to *V*. **d**, previously published genetic mapping of *V* vs *v*. **e**, candidate gene list within the *V* genetic interval with *Psat05G0805200* indicated in orange. **f**, Allelic/haplotype variation across the diversity panel for *Psat05G0805200* showing the cluster of parchmentless accessions in Hap2. **g**, summary and distribution of haplotypes of *P* and *V* among the parchmentless accessions.

**Extended Data Fig. 8.**
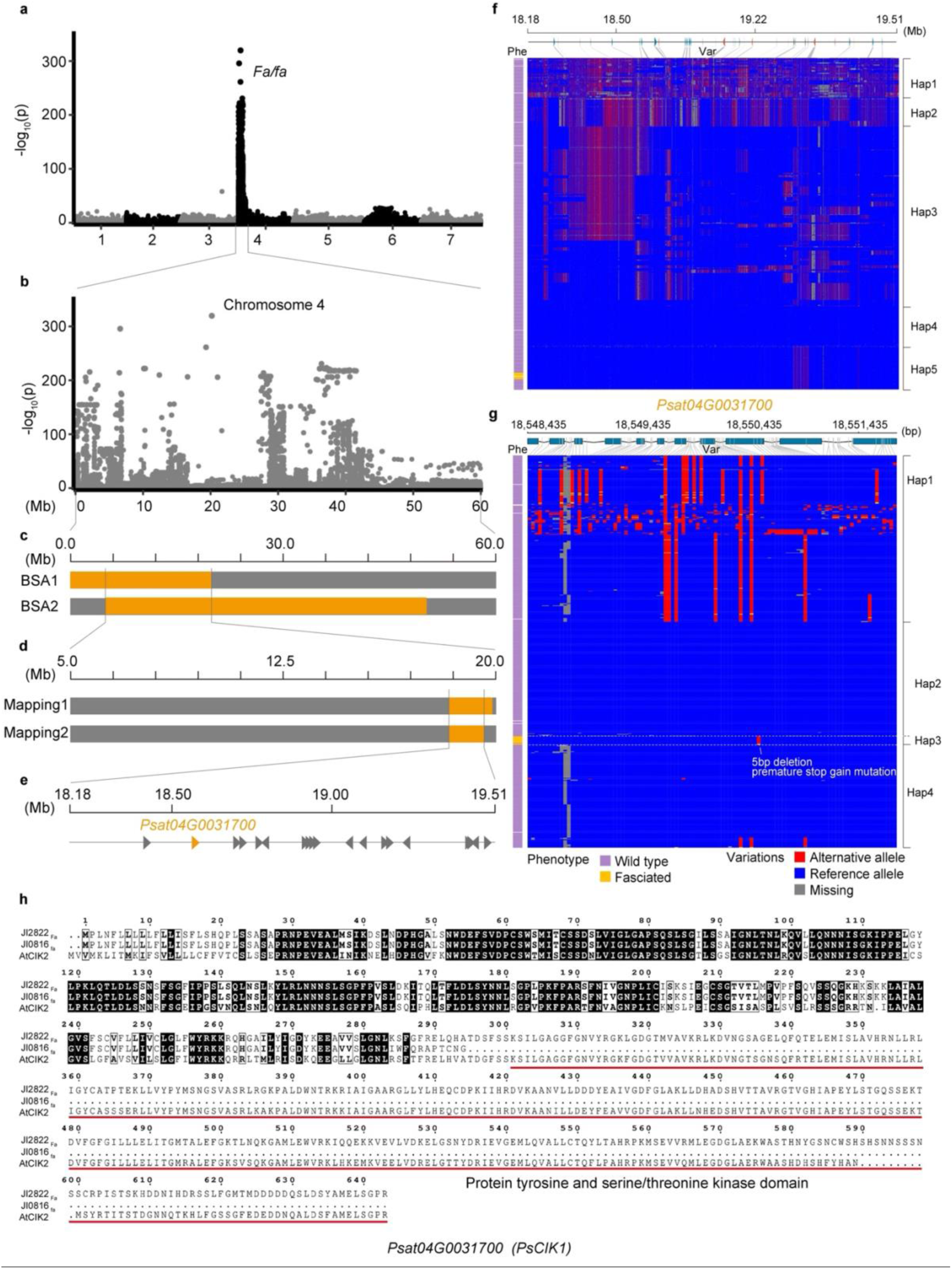
Gene and allele discovery of gene candidate *PsCIK1* for *Fa*. **a,** Manhattan plot of GWAS based on the ZW6 genome reference for fasciation revealing a peak of significance between 0 and 40 Mb on chromosome 4; **b,** Close up of Manhattan plot of GWAS in the region of the peak in a; **c,** Bulked segregant mapping analyses in the F2 of the cross Caméor (*FaFa*) x JI0814 (*fafa*), and in the F2 of JI2822 (*FaFa*) x JI0816 (*fafa*), further narrowed down the genetic interval; **d,** Fine mapping in Caméor x JI0814 (Mapping 1), with 8 pairs of KASP markers narrowed the *Fa* region to chr4: 18144306-19945776 (Supplementary Table 25); Fine genetic mapping in the JI0816xJI2822 population (Mapping 2) limited *Fa* to the interval chr4:18180969-19506907 (marker interval AX-183636277-AX183633456, Supplementary Table 17). **e,** Local detail of the genomic interval in panel d showing 20 protein-coding genes annotated as indicated. *Psat04G0031700* which encodes a Senescence-Associated Receptor-Like Kinase is highlighted in orange; **f,** a population-based haplotype clustering analysis across the diversity panel for the 1.33Mb *Fa* region identified showing a cluster of fasciated accessions in Hap5; **g,** a population-based haplotype clustering analysis of *Psat04G0031700* (*PsCIK1*) showing a 5bp deletion associated with the fasciated phenotype, and all the fasciated accessions clustered into Hap3. **h**, Amino acid sequence alignment of *CIK1* proteins from the wild-type line (JI2822, *Fa*, *Psat04G0031700*), the mutant line (JI0816, *fa*, *Psat04G0031700*-5bp), and the ortholog from Arabidopsis (*AT2G23950.1, AtCIK2*).

**Extended Data Fig. 9.**
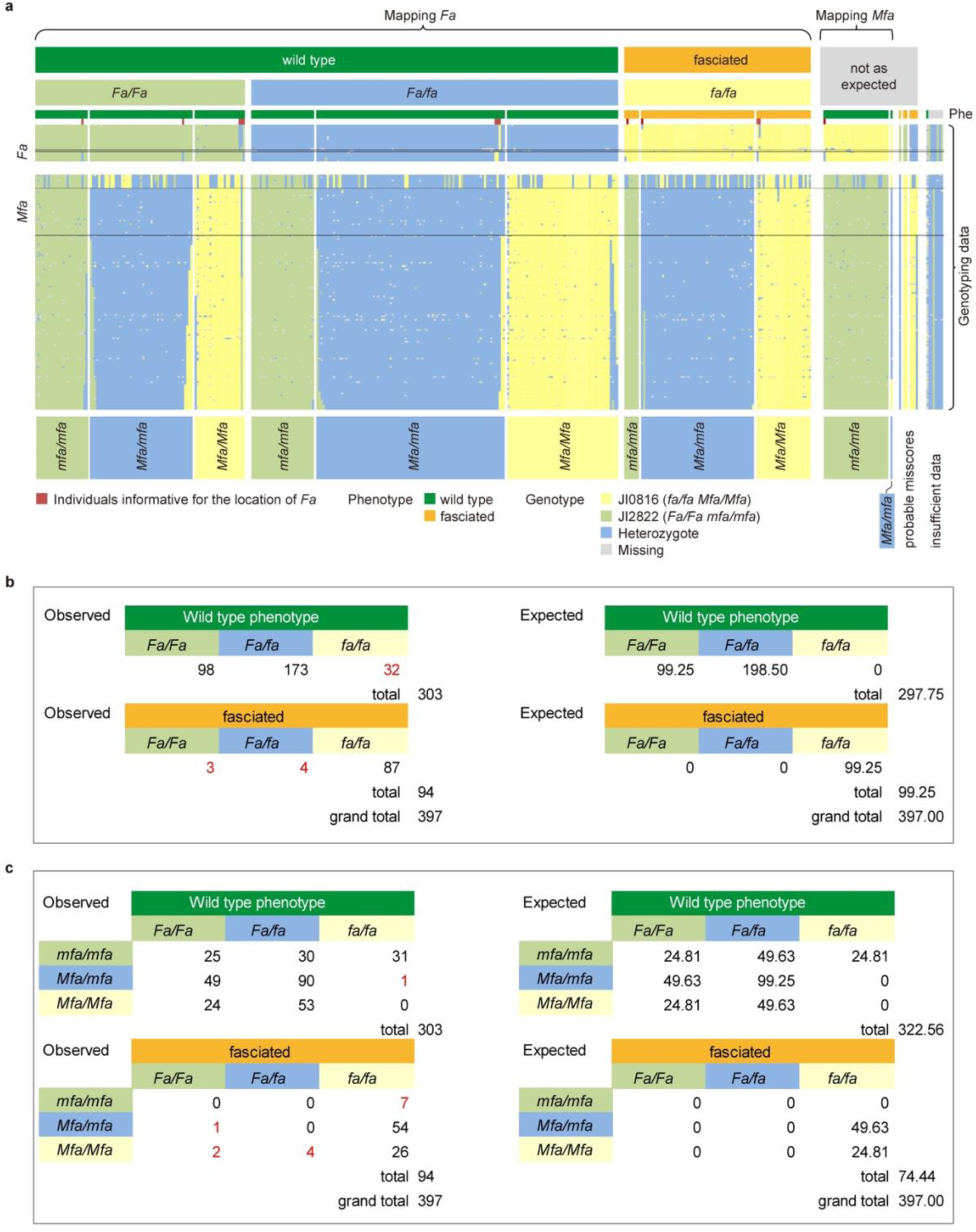
Segregation of *Fa* and *Mfa*. **a**, An excel spreadsheet is shown with genotype data (rows) for individuals in the JI0816xJI2822 F2 population (columns). The F2 individuals are sorted left to right according to their phenotype and their genotypic scores at *Fa* and *Mfa*. In the central upper part of the figure, homozygous JI0816 genotypes (*fafa*) are represented in yellow, homozygous JI2822 genotypes (*FaFa*) are represented in green, and heterozygotes (*Fafa*) are represented in blue. In the central lower part of the figure, homozygous JI0816 genotypes (*MfaMfa*) are represented in yellow, homozygous JI2822 genotypes (*mfamfa*) are represented in green, and heterozygotes (*Mfamfa*) are represented in blue. The limits of recombination intervals are marked by horizontal black lines. Wild-type (dark green) and fasciated (orange) phenotype scores are shown above the genotyping data. Homozygous and heterozygous genotypes at a proposed modifier locus, *mfa*, are shown below the genotyping data. F2 individuals informative for the positioning of *Fa* are marked with a red box; **b**, Tables explaining a one gene model of the summarised numerical data from panel a, where genotype *fa/fa* is fasciated; **c,** Tables explaining a two gene model of the summarised numerical data from panel a, showing postulated *Fa Mfa* interaction, where the dominant allele *Mfa* is required for fasciation to occur. In this model *fa/fa mfa/mfa* is wild type but *fa/fa Mfa/_* is fasciated. In both tables the numbers in red are F2 individuals with unexpected genotype/phenotype combinations.

**Extended Data Fig. 10.**
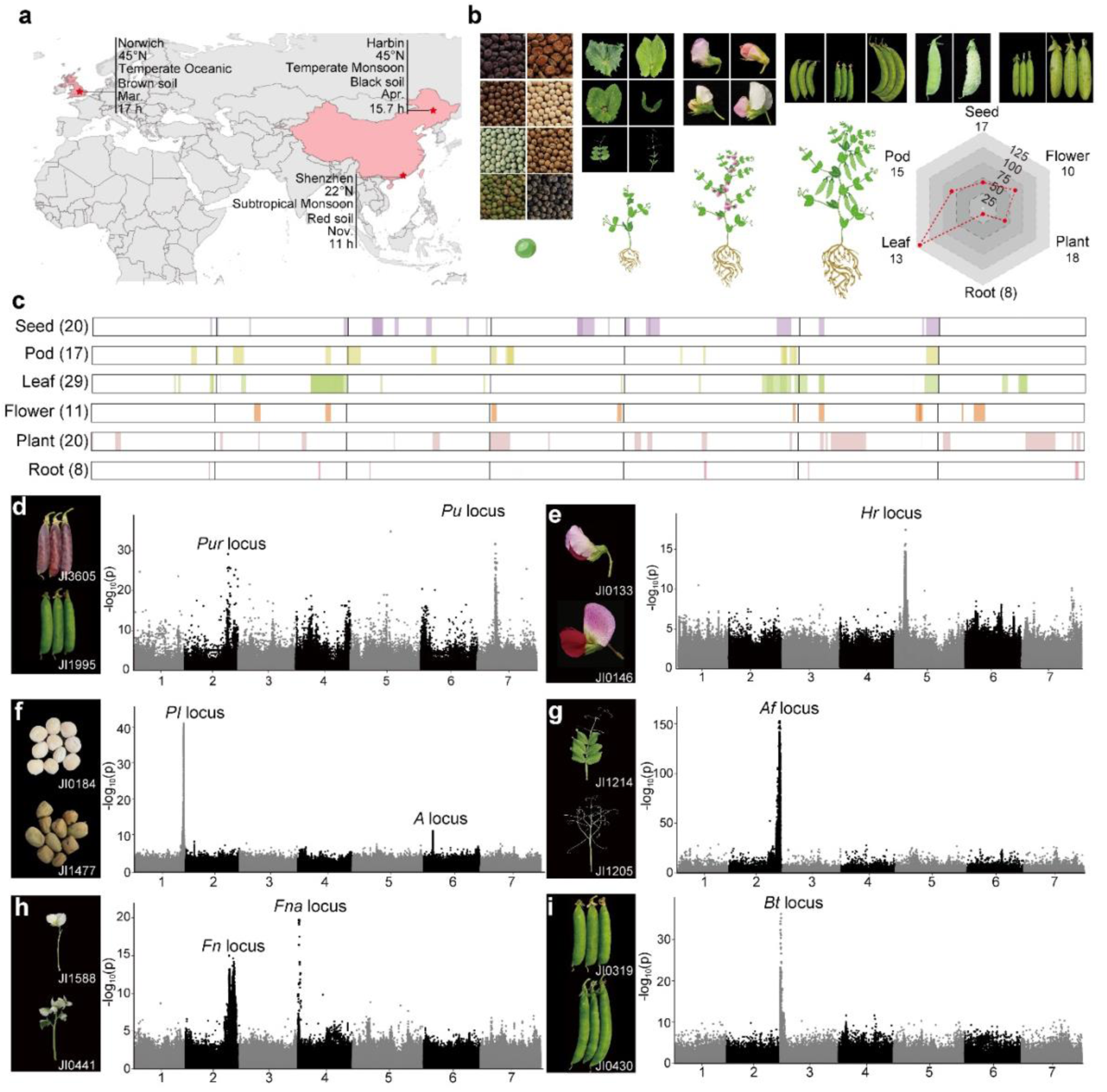
Identification of genomic loci conferring major agronomic traits. **a,** Multi-site phenotyping experiments were conducted to measure 81 traits from distinct climate zones, in three different locations: Southern China (22°N, Shenzhen), Northern China (45°N, Harbin), and the UK (52°N, Norwich). **b,** Illustrative photographs and drawings of phenotypic data collected for four (seed, leaf, flower, pod) out of the six high-level trait categories (including plant architecture, and root) scored in this study. The points in the hexagon indicate the total number of sub-traits collected. The red line indicates the total number of phenotypes collected (Supplementary Table 27). **c,** Significant marker-trait associations (MTAs) with genetic effects for component traits from seeds, pods, leaves, flowers, roots and plant architecture. The total number of sub-traits for each category is shown in parentheses. Genomic locations for the MTAs are given along the seven chromosomes of pea; **d**, Manhattan plot of GWAS based on the ZW6 genome reference, explaining green pod vs purple pod phenotypes corresponding to the *Pur* and *Pu* loci **e,** Manhattan plot of GWAS based on the ZW6 genome reference, explaining variation in flowering time corresponding to the *Hr*^85^ locus; **f,** Manhattan plot of GWAS based on the ZW6 genome reference, explaining brown vs black hilum colour phenotypes, corresponding to the *Pl* locus^53^ **g,** Manhattan plot of GWAS based on the ZW6 genome reference, explaining the leaf with leaflets and tendrils vs leaf with tendrils only phenotypes, corresponding to the *Af* locus^54^ **h,** Manhattan plot of GWAS based on the ZW6 genome reference, explaining variation in flower number corresponding to the *Fn* and *Fna* loci. **i,** Manhattan plot of GWAS based on the ZW6 genome reference, explaining the acute vs blunt pod tip phenotypes corresponding to the *Bt* locus

## Notes

### Competing Interest Statement

The authors have declared no competing interest.

